# Growth-coupled microbial biosynthesis of the animal pigment xanthommatin

**DOI:** 10.1101/2024.10.04.616593

**Authors:** Leah B. Bushin, Tobias B. Alter, María V.G. Alván-Vargas, Lara Dürr, Elina C. Olson, Mariah J. Avila, Òscar Puiggené, Taehwan Kim, Leila F. Deravi, Adam M. Feist, Pablo I. Nikel, Bradley S. Moore

## Abstract

The mining of genomes across life has unearthed a bounty of biosynthetic potential to diverse molecules key to a biobased future. While the heterologous expression of metabolic pathways has achieved broad success, most approaches suffer a similar fate in low initial production levels that require extensive, resource-heavy iterative strain engineering refinement. Herein we introduce a growth-coupled biosynthetic (GrowBio) strategy that irrevocably connects microbial growth with specialized compound production. We demonstrate the plug-and-play versatility of GrowBio in the production of the structurally complex animal biopigment xanthommatin, a color-changing ommochrome with material and cosmetic potential. Xanthommatin biosynthesis directly fuels growth of a newly designed *Pseudomonas putida* 5,10-methylenetetrahydrofolate auxotroph (PUMA). Aided by genome-scale metabolic modeling, PUMA was designed and built to be controlled by endogenous formate co-produced as a coupled biosynthetic byproduct in the multistep conversion of tryptophan to xanthommatin. Adaptive laboratory evolution was utilized to streamline xanthommatin’s gram-scale bioproduction via growth rate selection, establishing GrowBio as a promising biotechnological approach for establishing and optimizing the microbial production of value-added molecules.

## Introduction

Xanthommatin (Xa) is a biopigment that distinguishes a select group of species from the animal kingdom for color patterning (monarch butterfly),^1^ camouflage (squid and octopus),^2^ nuptial coloration (dragonfly),^3^ and compound eye vision (housefly) (**Fig. 1a**).^4^ Although common in arthropods and cephalopods, this biochrome is unusual among natural pigments due its ability to change colors in the presence of multispectral light, biochemical redox agents, and electrochemical potentials.^5^ These characteristic features have inspired numerous studies to understand the origin and function of xanthommatin in animals as well as to synthesize and test it as a broad-spectrum electrochromic, antioxidant material. Recently, xanthommatin has been explored in applications ranging from color-changing displays,^6^ coatings,^7^ dyes,^8^ and UV-protectants.^5,9,10^ Despite this promise, xanthommatin has yet to be produced biosynthetically in a cell factory. Such an achievement could both provide a sustainable production method and enable the creation of engineered living coloration.

**Figure 1:**
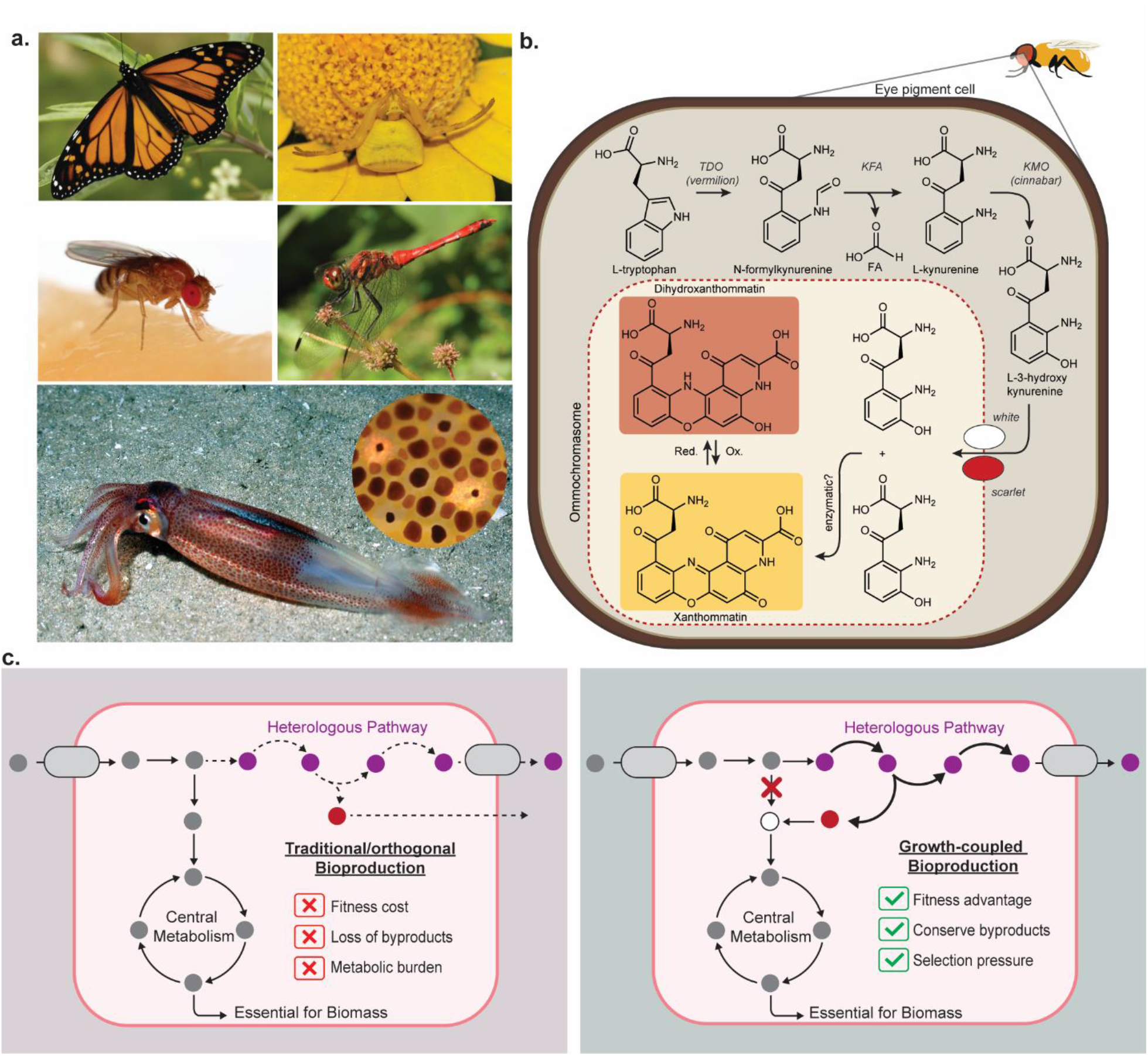
**a**. Examples of arthropods and cephalopods that produce ommochrome visual pigments such as xanthommatin. Photo credits: *(clockwise from top left) Danaus Plexippus*, Armon, CC BY-SA 3.0; *Thomisus onustus*, Alvesgaspar, CC BY-SA 3.0; *Sympetrum darwinianum*, TekuraDF, CC0; *Doryteuthis pealeii*, Andrew David, CC BY 2.0; Inset, Minette, CC BY 2.0; *Drosophila melanogaster*. Sanjay Acharya, CC BY-SA 4.0; **b**. The xanthommatin biosynthetic pathway in *Drosophila melanogaster* where formic acid (FA) is produced as a byproduct. 3HK is incorporated into pigment granules (called ommochromasomes) by a heterodimer composed of ABC transporters Scarlet and White. **c**. Schematic comparison of traditional (i.e., orthogonal left) versus growth-coupled (right) heterologous production designs. In the growth-coupled model, an auxotrophy is engineered for complementation with a co-produced heterologous metabolic byproduct (red circle) that directly connects essential biomass generation with product biosynthesis. Dashed arrows in the traditional design represent low activity through the heterologous pathway. Thick arrows in the growth-coupled design represent high(er) activity through the heterologous pathway.

In recent years, the production of numerous microbial pigments like carotenoids, quinones, and melanins^11–16^ has been engineered in synthetic cells as a replacement strategy for synthetic colorants. Xanthommatin is distinctive for its origin from the condensation of two molecules of L-3-hydroxykynurenine (3-HK) that derive from L-tryptophan via a 3-step oxidative pathway (**Fig. 1b)**. While the biochemical pathway to 3-hydroxykynurenine is well established across unicellular and multicellular life, the distribution of xanthommatin is largely limited to insects, arachnids, and cephalopods, as well as humans,^17^ suggesting that the dimerizing process is similarly rare. While the biosynthetic dimerization mechanism remains unknown to the pyridino-phenoxazine moiety, numerous reports have identified enzymatic and autoxidative mechanisms. For example, nonenzymatic oxidation has been described in human eye lenses, while the enzymes phenoxazinone synthase, *cardinal* peroxidase, and tyrosinase have been implicated in insects.^18,19^ Given the uncertainty of the biosynthetic coupling process, we set out to achieve a high titer bioproduction of the 3-hydroxykynurenine monomer in an appropriate microbial cell factory.

The heterologous expression of biosynthetic pathways has achieved broad success in the microbial production of diverse compounds across life, including new-to-nature molecules like pharmaceuticals, yet most expression strategies suffer a similar fate in initial low production levels.^20^ Metabolic engineering workflows, which rely on iterative design-build-test-learn cycles, are powerful solutions to optimize production rates, yields, and titers, yet are labor, resource, and time intensive (**Fig. 1c**). Growth-coupled bioproduction, in which the production of a desired molecule is intimately linked to the growth of the producing strain, offers an underutilized metabolic engineering strategy to rapidly overproduce a heterologous product^21–24^ such as xanthommatin. While growth-coupled bioproduction approaches have typically achieved increases in production of simple primary metabolites^25^ by rewiring new metabolic pathways to increase flux through target enzymes or pathways, we sought to develop a new strategy whereupon we could directly couple biosynthesis and cell viability in a portable, plug-and-play methodology based on one-carbon (C1) metabolic recapture.

The growth-coupled biosynthetic process to heterologous products takes advantage of the general concept that most biosynthetic pathways are not atom efficient and often co-produce carbon-based byproducts en route to the ultimate desired metabolic product. We reasoned that if we could rewire a cell’s metabolism to become reliant upon a heterologously produced co-product, such as a C1 metabolite, then we could fully connect growth to bioproduction through one-to-one coupling. As a proof of principle to evaluate the bioproduction potential of the growth-coupled biosynthetic (GrowBio) approach to sustainable products, we report here the creation of an engineered *Pseudomonas putida* strain fully dependent on formate, endogenously produced during the biosynthesis of the colorful animal pigment xanthommatin. A stepwise metabolic engineering approach resulted in the enforced production of xanthommatin at the gram/liter level (a first-case example in either natural or synthetic organisms) with minimal optimization, thereby showcasing the power, ease, and efficiency of the GrowBio approach to complex specialized metabolites.

## Results

### Rationale for linking C1 metabolism to the GrowBio workflow and selection of *P. putida* as a host strain

The most common carbon-based biosynthetic byproducts are C1 molecules, such as CO_2_, formate, and formaldehyde, produced during wide-ranging enzymatic reactions common in cellular metabolism. These C1 molecules are generally secreted or poorly assimilated back into central metabolism. In recent years, seminal progress has been made towards engineering model, industrially relevant microorganisms for growth on these very same C1 compounds as sustainable feedstocks.^26–31^ This has been accomplished by rewiring metabolism to introduce synthetic auxotrophies that are relieved by assimilation of exogenous C1 compounds whereupon they enter central metabolism via a series of interconnecting metabolic pathways via methionine and folate cycles. These essential cycles support purine and thymidine synthesis and the remethylation of homocysteine.

To explore the bioengineering potential of a GrowBio strategy linked to C1 metabolism, we first designed a platform strain that could be broadly employed irrespective of the product as long as an appropriate C1 byproduct is released. We selected the industrial strain *P. putida* as a bacterial chassis for its versatile metabolism and high tolerance to toxic substrates and products,^32^ paired with resistance to oxidative stress as 3-hydroxykynurenine mediates the production of reactive oxygen species.^33^ With respect to our goal to specifically overexpress xanthommatin, three additional features stood out. First, xanthommatin is toxic to many microbial species, both bacteria and yeast, but *Pseudomonas* species are tolerant.^34^ Second, pseudomonads encode the tryptophan-to-kynurenine pathway and can serve as an ideal gene source. And third, although the oxidative dimerization of 3-hydroxykynurenine to xanthommatin has not been firmly established in animals, some *Pseudomonas* species produce cinnabarinic acid, which contains a related phenoxazine core moiety formed from a similar condensation of two molecules of 3-hydroxyanthranilic acid.^35^

### Computational design and construction of *P. putida* 5,10-methylenetetrahydrofolate auxotroph (PUMA)

We engineered a synthetic formate auxotrophy in *P. putida*, which was related to a recent design in *Escherichia coli*.^36^ Using genome-reduced *P. putida* EM42 as the parent strain, a derivative of KT2440 optimized for heterologous gene expression,^37^ we targeted the biosynthetic reconfiguration of the essential cofactor 5,10-methylenetetrahydrofolate (MTHF). In *P. putida*, MTHF is formed by the transfer of C1 units to THF from serine and glycine via serine hydroxymethyltransferase (SHMT, encoded by *glyA-I* and *-II*) and the glycine cleavage system (encoded by *gcvTHP-I* and *-II*), respectively (**Fig. 2**). Moreover, MTHF is also a precursor for 10-formyl-THF and purine biosynthesis in *P. putida*. To restore MTHF biosynthesis, we employed the formate assimilation pathway from *Methylobacterium extorquens*^38^ that transfers formate directly to THF before being converted to MTHF.

**Figure 2:**
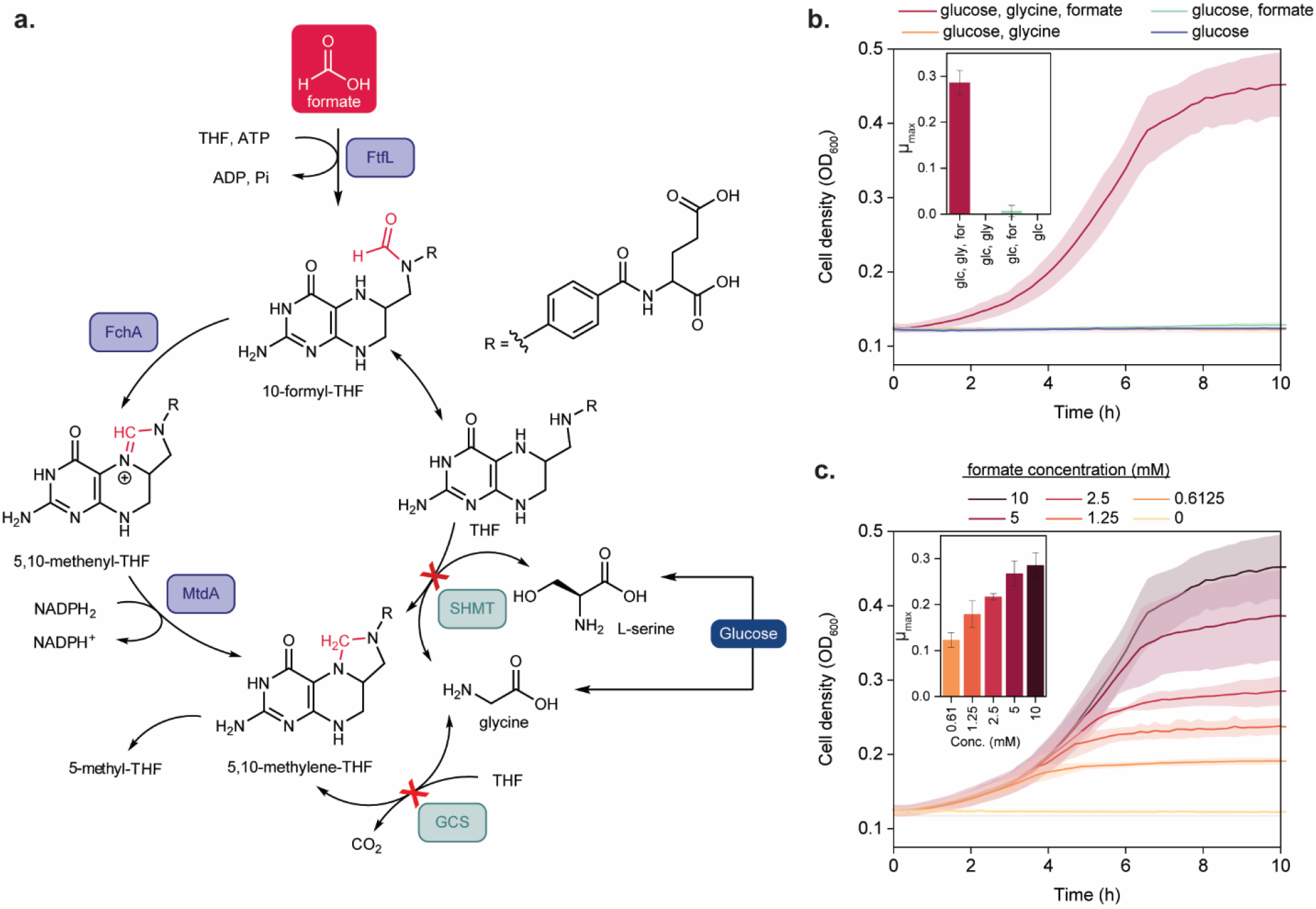
Design and characterization of the *P. putida* MTHF auxotroph PUMA. **a**. The MTHF pathway in *P. putida* involves the transfer of C1 units to THF from serine and glycine, both of which were blocked as shown by the red crosses. Addition of the formate assimilation pathway via FtfL, FchA, and MtdA (in purple) restores MTHF biosynthesis from formate; **b**. A growth plot of the PUMA strain in de Bont minimal (DBM) medium containing glucose (20 mM) in the absence or presence of formate (10 mM) and glycine (10 mM) along with an inset showing growth rate (*µ*_max_). Growth of PUMA required glucose (glc), glycine (gly), and formate (for); **c**. A growth plot of the PUMA strain in DBM containing glucose (20 mM), glycine (10 mM) and formate at the concentrations indicated along with an inset showing growth rate (*µ*_max_, h^-1^) at each formate concentration. Fitness of PUMA was dependent upon increased concentrations (conc.) of formate. Average values and standard deviation of three biological replicates are presented.

We thus interrogated the formate auxotrophy design *in silico* using the genome-scale metabolic model *i*JN1463 for *P. putida*^39^ and the growth-coupling algorithm gcOpt^22^ implemented in the Growth-Coupling-Suite^40^ that predicted deleting *gcvTHP* and *glyA* should establish the auxotrophy (**Fig. S1**). Calculation of the 2D flux space spanned by the growth rate and formate assimilation using the metabolic model of the double-deletion strain shows a strict formate dependence, which corresponds to a positive lower formate uptake bound for any given growth state (**Fig. S1a**). The respective metabolic flux rearrangements caused by the design direct THF-activated formate towards purine biosynthesis via inosine 5-monophosphate (IMP, 87% of assimilated formate), as well as L-methionine (12%) and pyrimidine (1%) biosynthesis via MTHF (**Fig. S1b**). The model also identified other genes that, in theory, could also mediate the synthesis of formyl-THF from formate, such as phosphoribosyl glycinamide formyltransferase (PurT) and THF-dependent phosphoribosylglycinamide formyltransferase (PurN). Given that the functions of these genes remain to be tested, we maintained both *purT/purN* in our initial strain design.

We built the *P. putida* synthetic MTHF auxotroph (PUMA) strain by deleting both copies of *glyA* and the two *gcvTHP* operons (**Fig. 2a**). We also inserted an optimized formate assimilation module, consisting of the genes encoding formate-THF ligase (FtfL, UniProt: Q83WS0), methenyl-THF cyclohydrolase (Fch, UniProt: Q49135), and methylene-THF dehydrogenase (MtdA, UniProt: P55818), into the *pha* locus, thereby deleting the native genes involved in polyhydroxyalkanoate biosynthesis^27,30^ that would otherwise act as a carbon sink. All genome modifications were carried out using I-SceI-assisted homologous recombination and, when necessary, CRISPR-Cas9 counterselection.^41–43^ Following construction, we tested growth of the PUMA strain in de Bont minimal (DBM)^44^ medium containing glucose, in the absence or presence of formate. The PUMA strain was, as predicted, unable to grow when glucose was included as the only carbon source. In the presence of glucose and exogenous formate, modest growth was detected. We suspected that endogenous levels of glycine due to the *glyA* deletion could be limiting due to the deletion of SHMT. However, simulations utilizing metabolic models of the wildtype and PUMA strains both were predicted to reach the same maximum growth rates without cofeeding glycine (**Fig. S1**), which suggested that the metabolic capacity to synthesize glycine was not impaired in PUMA and rather points to an unfavorable regulatory regime. When exogenous glycine was included in the medium with glucose and formate, robust growth was restored (**Fig. 2b**). Moreover, specific growth rate and maximum cell density achieved were dependent upon formate concentration. An 8-fold increase in concentration of supplemental formate (from 0.6125 mM to 10 mM) resulted in a 2.3-fold increase in both the growth rate and maximum cell density (**Fig. 2c**).

### Growth-coupled biosynthesis with endogenous formate from the kynurenine pathway

To test whether endogenously generated formate could also rescue growth and simultaneously empower heterologous biosynthesis, we selected the kynurenine pathway, in which the carbon atom at position C-2 of tryptophan is oxidatively removed as formate (**Fig. 3a**). The kynurenine pathway leads to diverse metabolic products with important roles in animals and is also active in pseudomonads. *P. aeruginosa* and *P. fluorescens*, for instance, produce anthranilate and the siderophore thioquinolobactin, respectively,^45–47^ from kynurenine. We sourced genes from the fluorescent pseudomonad *Pseudomonas* sp. DTU12.1, which harbors an active thioquinolobactin biosynthetic gene cluster (*qbs*),^48^ to construct two plasmid-based pathways for the synthesis of kynurenine and 3-hydroxykynurenine (**Fig. 3b**).

**Figure 3:**
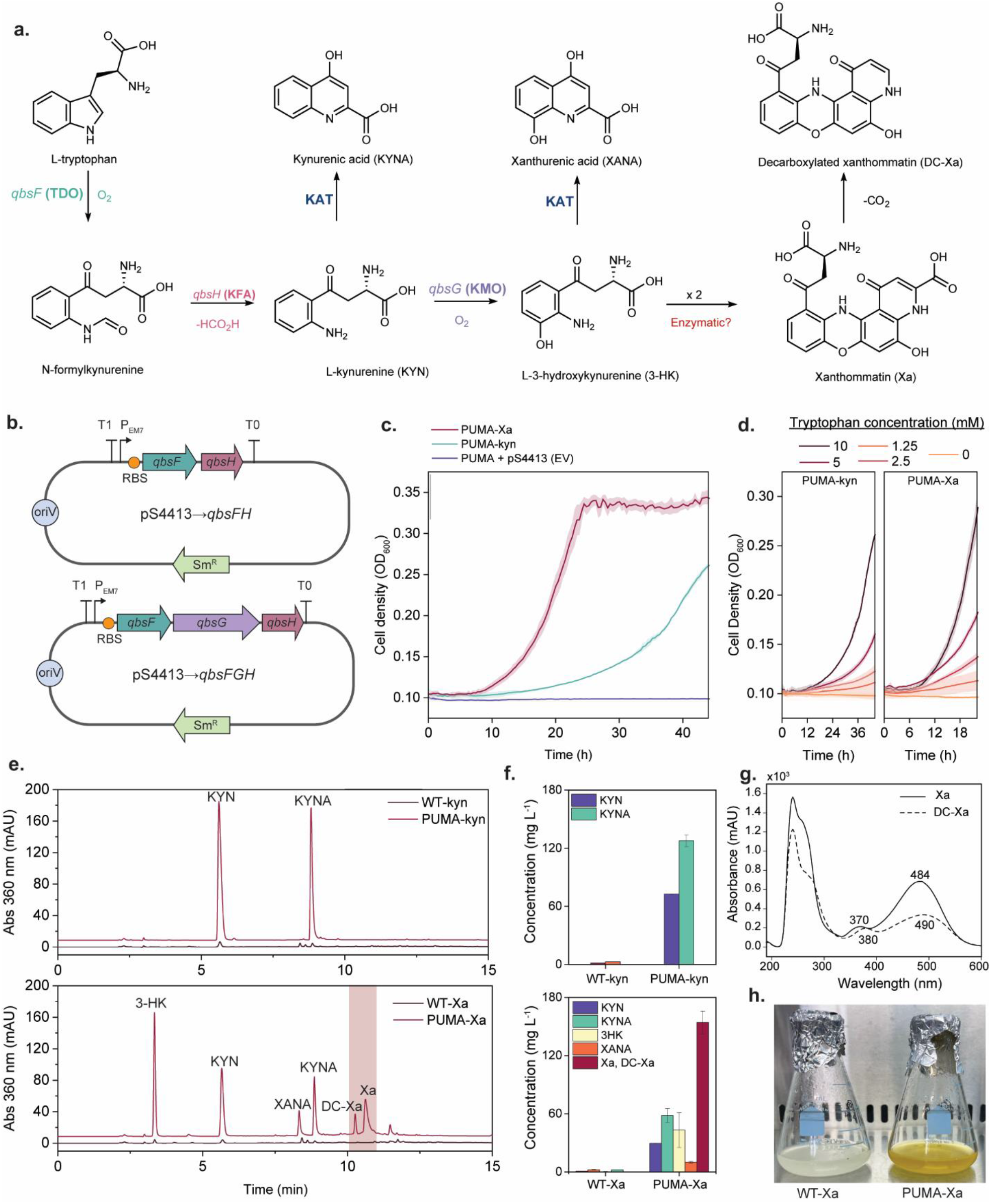
Growth-coupled biosynthesis of kynurenine products in PUMA. **a**. Three-step biosynthetic pathway converting tryptophan to 3-hydroxykynurenine (3HK) with formate (-HCO2H) as a co-produced byproduct. Kynurenine and 3HK are further converted to kynurenic acid, xanthurenic acid, and xanthommatin; **b**. Design of the kynurenine (KYN) and 3HK expression plasmids; **c**. Tryptophan supports the growth of *P. putida* strains PUMA-kyn and PUMA-Xa, but not PUMA, in the absence of formate; **d**. Growth rate and maximum cell density of PUMA-kyn and PUMA-Xa increase with higher concentration of L-Trp; **e**. Reversed-phase HPLC analysis of *P. putida* strains WT-kyn, PUMA-kyn, WT-Xa, and PUMA-Xa and the production of 3HK, KYN, xanthurenic acid (XANA), kynurenic acid (KYNA), xanthommatin (Xa), and decarboxylated Xa; **f**. Metabolite production titers of WT-kyn, PUMA-kyn, WT-Xa, and PUMA-Xa with 5 mM supplemental L-Trp; **g**. UV-vis spectra of Xa and DC-Xa crude mixture; **h**. Culture flasks of WT-Xa and PUMA-Xa.

The simplest construct consisted of two genes, *qbsF* and *qbsH*, that encode tryptophan 2,3-dioxygenase (TDO) and kynurenine formamidase (KFA), respectively, to convert tryptophan to *N*-formylkynurenine and ultimately kynurenine plus formate (**Fig. 3b**). Upon introduction of *qbsFH* into the formate auxotroph to create strain PUMA-kyn (LBB06), we evaluated its growth via a series of selective conditions. We observed that tryptophan could replace formate to sustain growth only in PUMA-kyn, but not PUMA (**Fig. 3c**), suggesting that MTHF biosynthesis was supported exclusively by the kynurenine pathway and not by native host pathways functioning as escape routes. Significantly, higher concentrations of exogenous tryptophan resulted in a faster growth rate, higher maximum cell density (**Fig. 3d**), and the overproduction of kynurenine and kynurenic acid, the latter of which is a likely product of an endogenous transaminase (**Fig. 3a,e**). We measured ∼45-fold increases in production of kynurenine and kynurenic acid in PUMA-kyn in comparison to the parent strain EM42 in which we similarly introduced the *qbsFH* plasmid (WT-kyn, LBB11) (**Fig. 3f, S2a**,**b**), underscoring the metabolic boost mediated by the growth-coupled approach.

We similarly constructed a three-gene pathway to 3-hydroxykynurenine that further included *qbsG* encoding kynurenine monooxygenase (KMO) (**Fig. 3b**). The resulting strain PUMA-Xa (LBB07) was also dependent upon tryptophan and glycine for growth (**Fig. 3c,d**). In this case, we observed the production of 3-hydroxykynurenine (43.1 mg/L) and its transaminated product xanthurenic acid (10.1 mg/L), in addition to kynurenine (29.5 mg/L) and kynurenic acid (58.1 mg/L) after 3 days of cultivation, each of which were produced at considerably greater titers than in EM42 expressing *qbsFGH* (WT-Xa, LBB12) (**Fig. 3e,f, S2c**,**d**). Significantly, we observed profound color changes from yellow to orange during the growth of PUMA-Xa, but curiously not in WT-Xa (**Fig. 3h**), that were not due to the kynurenine-based monomers. HPLC-MS analysis revealed the formation of two new compounds in the PUMA-Xa supernatant with absorption spectra (**Fig. 3d**) and masses consistent with the 3-hydroxykynurenine-based dimers xanthommatin (Xa) ([M+H]^1+^_obs_ 424.0786, [M+H]^1+^_calc_ 424.0776) and decarboxylated xanthommatin (DC-Xa) ([M+H]^1+^_obs_ 380.0882, [M+H]^1+^_calc_ 380.0878) (**Fig. S3**) at approximately 154.3 mg/L (**Fig. S2c**,**d**). We isolated DC-Xa and confirmed its identity by one- and two-dimensional NMR (**Fig. S4**).

### Adaptive laboratory evolution (ALE) to optimize growth-coupled biosynthesis

Despite the metabolic advantage of strains PUMA-kyn and PUMA-Xa in producing kynurenine molecules, these strains require supplemental glycine and tryptophan for growth even though the cells have endogenous routes to synthesize these amino acids from glucose. We thus applied adaptive laboratory evolution (ALE) to wean the system off these supplemental amino acids.^49^ We first subjected the formate auxotroph PUMA to ALE to support growth in the absence of exogenously supplied glycine using a controlled weaning and selection protocol on an automated platform.^50^ Surprisingly, all eight independent replicate ALE experiments using an independent isolate of the same glycine-requiring starting PUMA strain showed sustained growth when passaged without glycine after propagation from a growth-limiting glycine culture. There was, however, a significant lag phase for the initial passages in glycine free media before robust repeated growth was established (**Fig. 4a, S5**). A single isolate was then obtained from five of the eight endpoint populations, its growth profiled **(Fig. 4c**), and subjected to whole genome sequencing.^51^ All five isolates contained missense mutations in the gene *metK*, which encodes S-adenosylmethionine synthase, yet at different sequence locations (**Fig. 4c)**. No other genomic mutations were consistently found in all strains, implicating the unexpected role of *metK* activity in glycine metabolism and revealing the unbiased power of growth-coupled ALE to enhance strain platforms not predicted by rational-engineering based approaches.

**Figure 4:**
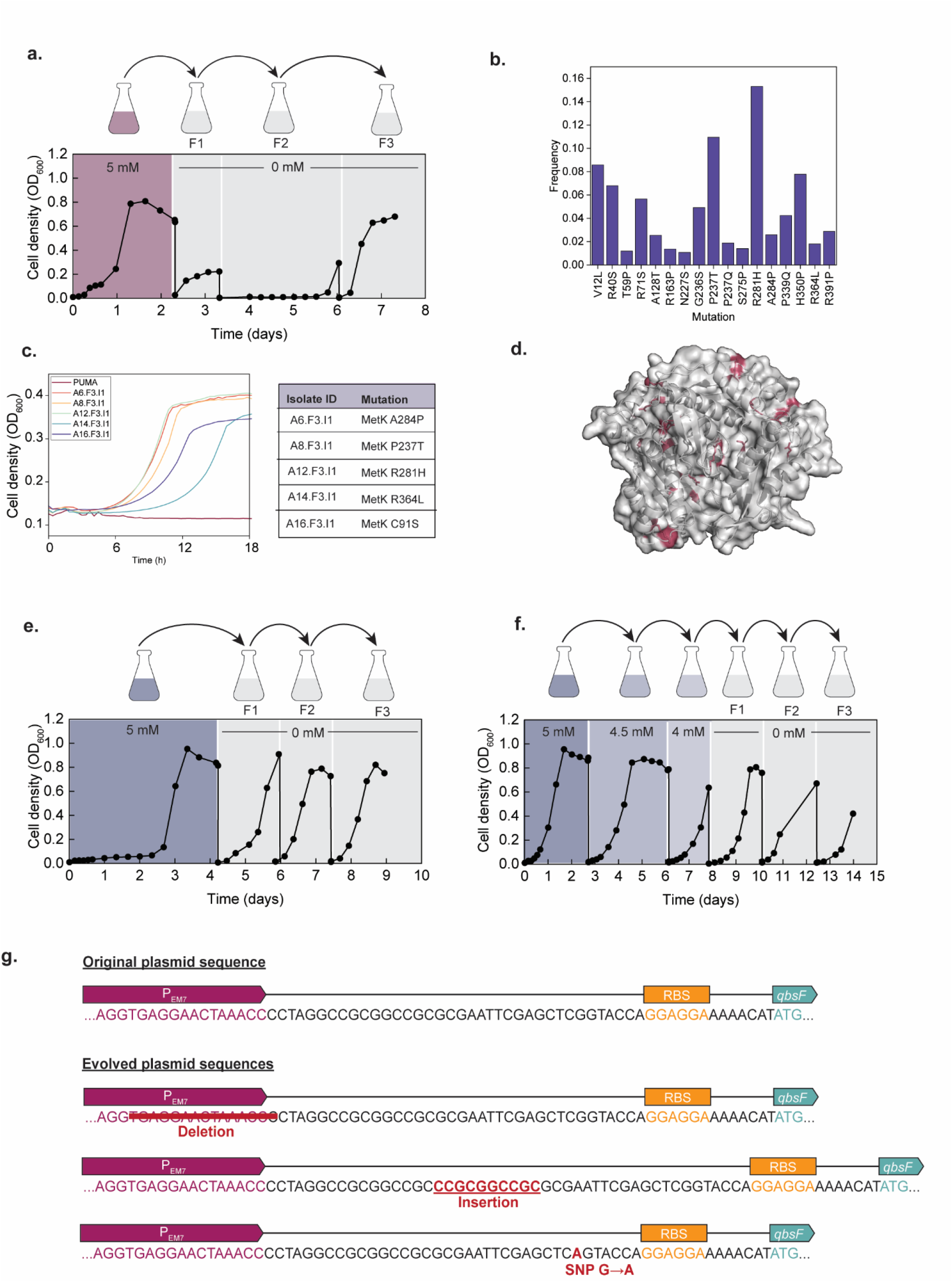
Adaptive laboratory evolution to select for a production strain utilizing glucose as the sole carbon source. **a**. A representative plot of growth during passaging to remove a glycine auxotrophy. Cells were grown in DBM medium containing glucose with supplementation of glycine as indicated; **b**. A plot of the mutational frequency for MetK mutations within an evolved population; **c**. A growth plot of evolved clones with various MetK mutations. Evolved clones, but not the parent strain, were able to grow in minimal media containing glucose and formate without glycine; **d**. Structural model of MetK with mutations highlighted; **e**. Representative passaging of the *qbsFH* tryptophan adaptation; **f**. Representative passaging of the *qbsFGH* tryptophan adaption; **g**. Representative sequences of the mutated promoted regions in plasmids isolated from evolved clones. Three varieties of mutations, including deletions, insertions, and single nucleotide polymorphisms, were observed. An example of each variety is shown.

To link the point mutations with the phenotypic observation of glycine weaning, we selected two MetK mutations, R281H and P237T, and engineered them into the PUMA base strain. In both cases, we observed glycine-independent growth comparable to the evolved mutant strains (**Fig. S6**), thereby confirming the role of MetK in relieving the glycine dependency. We next tested the prevalence and diversity of MetK mutants within a single population (labeled A8-F3-I0), which revealed a large number of *metK* mutations likely associated to many different subpopulations present from the short-term weaning ALE experiment (**Fig. 4b**). A library of *metK* amplicons was generated by PCR and subjected to ONT sequencing. Each amino acid position was then examined for the percentage of amplicons containing a certain mutation at that position. Seventeen mutations were found at a frequency of greater than 1%, with the top three most frequent being R281, P237T, and H350P. All five mutations found in the original single isolates, except for C91S, were also detected in the population analysis (**Fig. 4c**).

Next, we transferred the kynurenine (*qbsFH*) and 3-hydroxykynurenine (*qbsFGH*) plasmids (**Fig. 3b**) into the glycine-independent, evolved population A8-F3-I0 (renamed ePUMA) to generate ePUMA-kyn and ePUMA-Xa, respectively. Utilizing the population allowed us to fully sample the *metK* sequence space and avoid bias towards a single *metK* mutant. We then carried out a second round of ALE to remove the tryptophan dependency following the same weaning protocol as with the glycine requirement. Five distinct ePUMA-kyn and a further five ePUMA-Xa transformants were selected at random to initiate parallel evolution experiments (**Fig. S7**). As we experienced with the glycine weaning experiment, a rapid loss of the tryptophan requirement for ePUMA-kyn with growth was observed early in the ALE experiment cultures that were tested without supplementation periodically (**Fig. 4e, S7**). Evolution to wean PUMA-Xa from tryptophan also occurred in a relatively modest number of generations, with replicates exhibiting growth in the first or third tryptophan-free flask (**Fig. 4f, S5**). Five isolated clones were tested from each of the endpoint populations. All showed growth and produced their expected kynurenine products when cultured in DBM medium containing solely glucose (20 mM) without amino acid supplementation (**Fig. S8**). The resulting mutant strains were named ePUMA-kyn(WIn) and ePUMA-Xa(WIn) to notate their tryptophan independence (WIn).

To evaluate the genomic changes in the evolved glycine/tryptophan-independent strains ePUMA-kyn(WIn) and ePUMA-Xa(WIn), we selected two clones per population for whole-genome sequencing. Strikingly, none of the apparent genomic mutations were common to all clones. The pathway plasmids, however, all contained mutations in the P_EM7_ promoter region, including deletions, insertional repeats and single nucleotide polymorphisms (**Fig. 4g, S9**). To test the consequence of the modified P_EM7_ sequences, we isolated plasmids from several of the evolved clones and introduced them directly into hosts PUMA and ePUMA (clone A8.F3.I1). The resulting strains were indeed able to grow in tryptophan-free minimal media, indicating that the modified promoters were sufficient to impart tryptophan-independent growth (**Fig. S10**).

### Fed-batch production and chemical analysis of biosynthetic xanthommatin

We conducted a time course study in shaken-flask cultures of strains ePUMA-kyn(WIn) and ePUMA-Xa(WIn), comparing production levels to WT-kyn and WT-Xa, respectively, with glucose as the sole carbon source. We measured enhanced production in the kynurenine/kynurenic acid-producing strain ePUMA-kyn(WIn) (59.8 mg/L kynurenine and 181.2 mg/L kynurenic acid) on par with unevolved PUMA-kyn that required supplemental glycine and tryptophan (**Fig. S11**). The 3-hydroxykynurenine producing strain ePUMA-Xa(WIn), on the other hand, exhibited a beneficial shift in its metabolite profile with a decrease in the amount of the shunt product kynurenic acid relative to the amount of L-3-hydroxykynurenine. Xanthommatin was produced by ePUMA-Xa(WIn) when grown with glucose as the sole carbon source (**Fig. S11, 5a**), but not by WT-Xa, as was the case even with tryptophan supplementation. The quantities of the other monomeric metabolites were also ∼10-30× greater in ePUMA-Xa(WIn) for kynurenine, kynurenic acid, hydroxykynurenine, and xanthurenic acid.

We next tested the production capacity of the evolved strain in a bioreactor in DBM medium under fed-batch conditions. We analyzed growth and product formation in a 250 mL bioreactor in which 34 grams of glucose were continuously fed to ePUMA-Xa(WIn) over 72 h to give a dense, maroon-colored culture broth (**Fig. S12)**. The supernatant was collected and treated with ascorbic acid to precipitate xanthommatin. We observed that reduced xanthommatin and it decarboxylated analog DC-Xa precipitate out of aqueous media, while the kynurenine-based monomers remained in solution, providing a facile purification route to 551 mg of crude xanthommatin powder from 230 mL of culture, with a calculated titer of 2.4 g/L.

To evaluate the optoelectronic properties of biosynthetic xanthommatin, we tested its performance via a series of analytical assessments and compared the results directly to a chemically synthesized standard. The UV-vis absorbance spectra indicated a 35 nm red-shifted visible band from 435 to 470 nm in the biosynthetic xanthommatin relative to the synthetic xanthommatin (**Fig. 5b**). Despite this difference, the redox-dependent color change inherent to xanthommatin was apparent in both preparations, which also compared favorably to our past reports of synthetic xanthommatin^52^ (**Fig. 5c**). These qualitative changes were further supported by the reversible electron transfer as confirmed from cyclic voltammetry (**Fig. 5d**). From these data, we extrapolated the optical energy and fundamental energy band gaps of each product (**Fig. 5f**), corresponding to the visible and UV spectral regions, respectively. These data revealed strong similarities between the biosynthetic and synthetic xanthommatin samples, highlighting sustainable production of this promising biomaterial.

**Figure 5.**
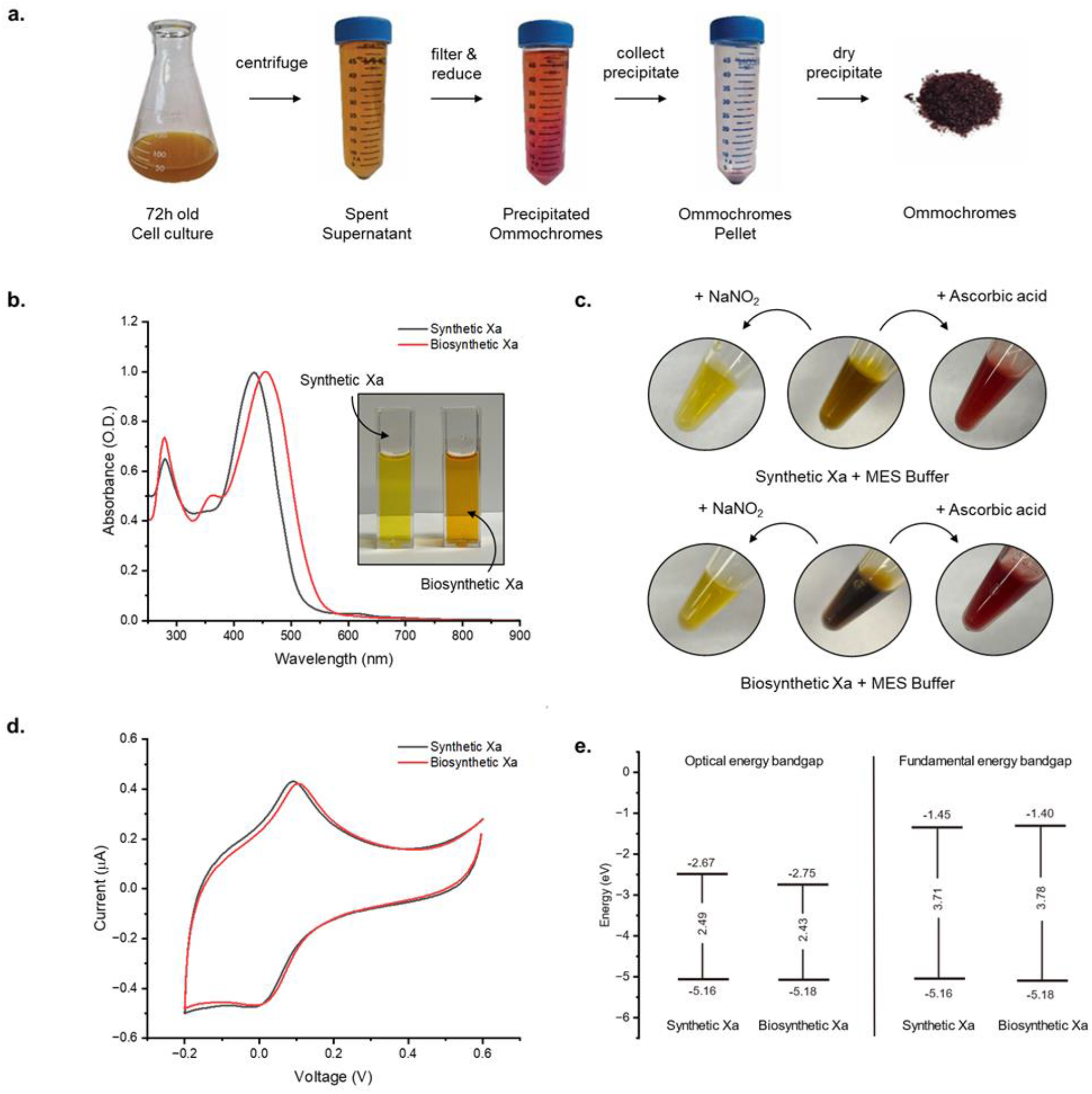
Analysis of optoelectronic properties of xanthommatin. **a**. Purification of xanthommatin ommochromes from shake flask cultures of ePUMA-Xa(WIn). Addition of ascorbate to aqueous supernatant causes precipitation of xanthommatin material. The other kynurenine-based monomers remain in solution; **b**. Comparison of UV-vis absorbance spectra in DMSO; **c**. Image of redox dependent color change of crude xanthommatin and biosynthesized xanthommatin of oxidized (yellow) and reduced (red) forms; **d**. Electrochemical analysis via cyclic voltammetry (CV) curves of each compound (0.5 mg/mL) in DMSO containing 0.1 M lithium triflate; **e**. Comparison of optical and fundamental energy bandgaps and associated HOMO/LUMO levels extrapolated from Tauc plots generated from the absorption data and values extrapolated from CV’s, respectively.

## Discussion

The microbial bioproduction of the colorful animal biopigment xanthommatin reduces to practice the metabolic engineering potential of the GrowBio strategy by intimately linking growth and compound production in an auxotrophic strain. By connecting the heterologous kynurenine pathway to C1 metabolism via an orthogonal formate assimilation pathway to MTHF in the PUMA auxotroph strain, we made the kynurenine pathway essential for primary growth via nucleic acid and methionine biosynthesis. While the never-before bioengineered xanthommatin was the target of this GrowBio campaign, we anticipate that this strategy is widely applicable to any biosynthetic product in which a C1 molecule is released during its construction. As such, this method has the potential to be agnostic to the product as long as it is not toxic and that formate (or a molecule that can be readily converted to formate) is produced and captured.

In the case of xanthommatin bioproduction via C1 growth-coupling, we learned several lessons, some of which were unexpected and others that have opened new lines of inquiry. We list five such lessons here. First, we showed that the PUMA base strain was reliant on glycine for growth, and upon ALE, genomics, and base editing, we identified and confirmed that seemingly random point mutations in the MetK S-adenosylmethionine synthase could cure the glycine growth requirement. This unexpected result gave rise to the evolved, glycine-independent strain ePUMA and will require additional inquiry to understand how *metK* mutations in *P. putida* connect with glycine biosynthesis. A second unexpected result was similarly illuminated in an ALE experiment when we weaned the ePUMA-Kyn and ePUMA-Xa production strains off exogenous tryptophan. In those cases, the relieving mutations were restricted to the expression plasmids, specifically to the P_EM7_ promoter region, ranging from deletions to insertional repeats that may regulate the demand on the tryptophan pool during initial growth.

The biosynthetic production of xanthommatin by PUMA-Xa was another unexpected outcome as we only initially expressed a three-gene cassette to produce 3-hydroxykynurenine. At that point in the engineering campaign, we had not yet tested one of the several oxidative dimerases identified in animals that support xanthommatin production. Since the condensation of 3-hydroxykynurenine has also been reported to occur autocatalytically, the production of xanthommatin in strain PUMA-Xa may similarly be enzyme independent or be the product of an as-yet identified enzyme in *P. putida* like catalase or laccase, which have been shown to catalyze the oxidative dimerization of *o*-aminophenols, like 3-hydroxyanthranilic acid to cinnabarinic acid.^53,54^ The production strain ePUMA-Xa(WIn) is nonetheless efficient in its production of xanthommatin with less than 50% of it substrate 3-hydroxykynurenine remaining in the fermentation at the time of harvesting. This unexpected conversion may portend ‘bonus’ enzyme discoveries in future GrowBio campaigns.

A fourth unforeseen outcome of the GrowBio experiment was our observation that PUMA-Xa achieved greater cell density more quickly and produced more kynurenine-based material than PUMA-Kyn. The only difference between these two constructs was the addition of kynurenine monooxygenase, which catalyzes the conversion of kynurenine to 3-hydroxykynurenine. As the preceding reaction by kynurenine formamidase releases formate to break the auxotrophy, we were pleased to observe that the addition of enzymes beyond the break not only benefit from the increased substrate pool, but also relieve the cell from product feedback^55^ that may reduce the efficiency of the deformylation process and thus growth. We anticipate that the GrowBio strategy will foster longer biosynthetic processes by pulling flux through the pathway.

The fifth benefit from the GrowBio process described here is pertinent to xanthommatin specifically. Not only does the growth-coupled strategy provide a practical production route to this promising, color-changing dye, but it also allows for the creation of engineered living coloration due to its high xanthommatin content. In recent years living biomaterials that contain live cells within polymeric materials have been engineered to respond to external stimuli.^56^ With xanthommatin’s optoelectronic properties, we are exploring its potential as a living material.

Beyond xanthommatin, the kynurenine pathway is a gateway to molecular diversity that may similarly benefit from growth-coupling. Such products include the flavoring agent methyl anthranilate, the antidepressant prodrug 4-chlorokynurenine,^57^ and a bevy of anthranilate-based bioactive alkaloids.^58^ The kynurenine pathway is just one of many that produce formate, or an equivalent C1 metabolite, as a co-produced byproduct of biosynthesis. As such, we anticipate that the C1 GrowBio strategy introduced herein will help accelerate the overproduction of new biosustainable materials.

## Data Availability

Genome sequencing raw files are available from the Sequence Read Archive with BioProject ID PRJNA1169116.

## Acknowledgments

Funding was generously provided by the National Institutes of Health (R01-GM085770 to B.S.M., F32-GM149120 to L.B.B.), the Office of Naval Research (N000142412649 to B.S.M. and L.F.D.), and the Novo Nordisk Foundation (NNF10CC1016517, NNF20CC0035580 and NNF18CC0033664 to P.I.N. and NNF20CC0035580 to T.B.A. and A.M.F.). We thank L. Jelsbak (DTU) for the kind gift of *Pseudomonas* sp. DTU12.1 and J. Turlin (DTU) and C. de Venecia (UCSD) for their research assistance.

## Author contributions

L.B.B., T.B.A., P.I.N. and B.S.M. conceived of the GrowBio strategy. L.B.B. conducted all the biochemical, metabolic, and genomic experiments, engineered the mutant strains with M.J.A. and O.P., and isolated xanthommatin with L.D. T.B.A. performed metabolic modeling simulations. L.B.B. and E.C.O. performed the ALE experiments under the supervision of A.M.F. L.B.B. and M.V.G.A.V. conducted the bioreactor experiments. T.K. and L.F.D. prepared and tested synthetic xanthommatin against biosynthetic xanthommatin. L.B.B. and B.S.M. wrote the first draft of the manuscript with subsequent input from all the authors. B.S.M. and P.I.N. supervised the project.

## Methods

### Bacterial strains and culture conditions

Bacterial strains employed in this study are listed in Supplementary Table S1. *E. coli* was grown at 37°C in LB medium. *P. putida* was grown at 30°C in LB medium for routine applications. Growth assays, bioproduction assays, and adaptive laboratory evolution (ALE) experiments were performed in de Bont minimal (DBM) medium (3.88 g L^-1^ K_2_HPO_4_, 1.63 g L^-1^ NaH_2_PO_4_, 2 g L^-1^ (NH_4_)_2_SO_4_), supplemented with trace elements (0.1 g L^-1^ MgCl_2_•6H_2_O, 10 mg L^-1^ ethylenediaminetetraacetic acid (EDTA), 2 mg L^-1^ ZnSO_4_•7H_2_O, 1 mg L^-1^ CaCl_2_•2H_2_O, 5 mg L^-1^ FeSO_4_•7H_2_O, 0.2 mg L^-1^ Na_2_MoO_4_•2H_2_O, 0.2 mg L^-1^ CuSO_4_•5H_2_O, 0.4 mg L^-1^ CoCl_2_•6H_2_O, 1 mg L^-1^ MnCl_2_•2H_2_O) and 3.6 g L^-1^ (20 mM) glucose. Glycine, formate, and L-tryptophan were added according to the strain requirements and experimental conditions at the concentrations indicated in the text. All liquid cultures were agitated at 250 r.p.m., unless otherwise stated. Solid culture media contained 15 g L^-1^ agar. Antibiotics were added as necessary at the following concentrations: 50 μg mL^-1^ kanamycin (Km), 10 μg mL^-1^ gentamicin (Gm) and 100 μg mL^-1^ streptomycin (Sm).

### Plasmids and cloning procedures

Plasmids and oligonucleotides used in this study are listed in Supplementary Tables 2 and 3, respectively. Oligonucleotides were purchased from Integrated DNA Technologies. Commercial kits and enzymes were used following the manufacturers’ protocols. PCR amplifications were performed using Phusion™ Hot Start II High-Fidelity DNA Polymerase (Thermo Scientific) or Q5® High-Fidelity DNA Polymerase (New England Biolabs (NEB)). FastDigest DpnI (Thermo Scientific) and BsaI (NEB) were used for restriction enzyme digests. T4 DNA Ligase (NEB) and T4 Polynucleotide Kinase (NEB) were used for ligation and phosphorylation, respectively. DNA assembly reactions were performed using NEBuilder® HiFi DNA Assembly Master Mix (NEB). Plasmid DNA was purified using the QIAprep Spin Miniprep Kit (Qiagen) and PCR amplicons were purified using either the QIAquick PCR Purification Kit (Qiagen) or the Zymoclean Gel DNA Recovery Kit (Zymo Research) following agarose gel electrophoresis. *E. coli* DH5α was employed as a general cloning host. *E. coli* DH5α λpir was used for cloning and maintaining plasmids that contain the conditional origin of replication R6K. Chemically competent *E. coli* cells were prepared using the calcium chloride method. Plasmids and assembled products were transformed into *E. coli* competent cells by heat-shock and plated onto LB agar containing the appropriate antibiotic. Colony PCR using OneTaq® Quick-Load® 2X Master Mix (NEB) was used to identify positive clones. *P. putida* was routinely transformed by electroporation using a 100 μL-aliquot of cells in a 2 mM cuvette with a voltage of 2.5 kV. Electrocompetent *P. putida* cells were prepared by washing with 0.3 M sucrose three times.

### Genome engineering of *Pseudomonas putida*

Genome modifications in *P. putida* were performed using I-SceI-mediated recombination, with the aid of CRISPR-Cas9 counterselection, as necessary, following established protocols. Briefly, to perform each modification, a pGNW2-derived suicide vector bearing homology regions corresponding to the target gene or locus was integrated into the chromosome. Co-integrants, confirmed by green fluorescence, were transformed with the vector pQURE6•H that encodes the I-SceI gene. When CRISPR/Cas9 counterselection was employed, a pS448•CsR derived vector carrying the appropriate target-specific guide RNA was co-transformed with pQURE6•H. Immediately following electro-shock, cells were recovered in liquid LB medium supplemented with 2 mM 3-methylbenzoic acid (3-*m*Bz) to induce I-SceI expression and 20 mM glycine to support growth of mutants. Within 3 h of recovery, the culture was plated on selective LB agar containing the appropriate antibiotic(s), 2 mM 3-*m*Bz, and 10 mM glycine. Plates were incubated until colonies were visible. Resolved colonies, confirmed by red fluorescence, were screened by colony PCR to test for the desired genotype. Correctly modified clones were serially passaged in antibiotic-free LB medium to cure pQURE6·H and the pS448·CsR counterselection plasmid, when present. Plasmid-cured clones were identified by the absence of red fluorescence and the loss of antibiotic resistance. During deletion of *gcv-II* LB medium was also supplemented with 10 mM formate to enrich for mutants. PCR fragments of *metK* mutants R281H and P237T were amplified from genomic DNA isolated from evolved clones and cloned into the pGNW2 suicide vector using the primers listed in Supplementary Table 3. During reengineering of *metK* mutations into the PUMA strain, LB agar was supplemented with appropriate antibiotic, 5 mM formate, and 5 mM glycine following transformation of pGNW2-derived vectors into the PUMA strain. Following transformation with the pQURE6·H vector, cells were recovered for 2-5 h in 1 mL LB supplemented with 2 mM 3-*m*Bz, 5 mM formate, and 5 mM glycine. All other steps in the reengineering followed the protocol described above

### Construction of CRISPR/Cas9 counterselection plasmids

Plasmids pS448·CsR_*glyA-II*_2 and pS448·CsR_*gcv-I*_2 were designed and constructed following an established protocol. Briefly, for each plasmid, the online tool CRISPy-web was used to design a suitable 20-nt spacer sequence with specificity for the genomic target. Two complementary oligonucleotides containing the spacer sequence and appropriate overhangs were phosphorylated and annealed to generate an insert for ligation between BsaI sites. The resulting insert and BsaI-digested linearized pS448·CsR vector were ligated, yielding a pS448·CsR derived plasmid encoding a sgRNA with specificity for the target locus. Primers sgRNA_*glyA-II*_2_F and sgRNA_*glyA-II*_2_R were used to make the insert with the spacer sequence targeting *glyA-I* and primers sgRNA_*gcv-I*_2_F and sgRNA_*gcv-I*_2_F to make the insert with the spacer sequence targeting *gcv-I*. Plasmids were verified by Sanger sequencing (Eurofins Genomics).

### Construction of pathway plasmids

Vector pSEVA4413 was PCR-amplified with primer pair pS4413_F_BamHI/pS4413_R_KpnI, to yield a linear backbone for cloning inserts between the BamHI and KpnI sites. The resulting PCR product was digested with DpnI restriction enzyme. The *qbsF* and *qbsH* genes were PCR-amplified from genomic DNA isolated from *Pseudomonas* sp. DTU12.1 with primer pairs qbsF_F/qbsF_R and qbsH_F/qbsF_R, respectively. A synthetic ribosomal binding site (RBS) (AGGAGGAAAAACAT) was introduced directly upstream of the *qbsF* start codon via the forward primer (qbsF_F). The PCR fragment containing *qbsH* also included the upstream *qbsG*-*qbsH* intergenic region which possesses a native RBS. All PCR fragments contained appropriate overhangs for DNA assembly. The *qbsF* and *qbsH* PCR fragments and the pS4413 backbone were joined using HiFi DNA assembly master mix, yielding pS4413-*qbsFH*. To construct pS4413-*qbsFGH*, the intact *qbsFGH* operon was directly amplified from *Pseudomonas* sp. DTU12.1 genomic DNA using the primer pair qbsF_F/qbsH_R. The resulting fragment, and the pS4413 backbone were joined using NEBuilder HiFi DNA assembly. Plasmids were verified by whole plasmid sequencing (Primordium Labs).

### Computational modeling

Strategies coupling L-3-hydroxykynurenine (3HK) synthesis to growth were computed using the Growth-Coupling Suite (GCS),^40^ a comprehensive framework based on the gcOpt algorithm.^22^ GCS utilizes constraint-based metabolic models and bilevel optimization to identify gene knockouts, heterologous knock-ins, and medium compositions that couple the activity of a specific reaction to cellular growth.

For this study, the genome-scale metabolic model *i*JN1463 of *Pseudomonas putida* KT2440 was used as the base model. To enable the production of 3HK as the xanthommatin precursor, a heterologous pathway was introduced to *i*JN1463, incorporating the reactions tryptophan 2,3-dioxygenase, kynurenine formamidase, and kynurenine monooxygenase. An exchange reaction for balancing the non-native product 3HK was also included, leading to the metabolic model *i*JN1463_3HK.

The GCS workflow was applied to *i*JN1463_3HK using the parameters and settings as described below. The kynurenine monooxygenase reaction was set as the objective function or target reaction to specifically compute strategies coupling the 3HK synthesis pathway to growth. To reduce the solution space, reactions whose associated genes are essential for *P. putida* or for which no associated genes exist were excluded as knockout targets. These included experimentally validated and model-predicted gene essentialities^39^, exchange, transport, and spontaneous reactions, as well as reactions assigned to essential model subsystems, i.e., Biomass and maintenance functions, Murein Recycling, Cell Envelope Biosynthesis, Iron uptake and metabolism, and tRNA Charging. A database of potential knock-in targets was derived from 28 genome-scale metabolic models representing different species, as described in detail^40^. To simultaneously optimize the minimal medium, a set of 25 substrates was defined from which GCS was allowed to select one as the sole carbon source per strategy. The maximum substrate uptake rate was uniformly constrained to 60 C-mmol gCDW^-1^ h^-1^. The GCS algorithm was run multiple times with different maximum numbers of genetic interventions to allow for a diverse set of growth-coupling strategies with experimentally tractable numbers of genetic interventions. The maximum number of knockouts and knock-ins was varied between two and five. The runtime of each GCS run was limited to 600 min. The final growth-coupling strategy, i.e., the knockout of the glycine cleavage system (GHMT2r) and serine hydroxymethyltransferase (GLYCL) using glucose as the sole carbon source, was selected from the accumulated set of solutions by manual inspection.

The MTHF auxotrophy enforced by the growth-coupling strategy and its relief by formate assimilation were confirmed separately. To represent the corresponding strain LBB05 the heterologous formate assimilation pathway from *Methylobacterium extorquens* was completed by adding the formate-tetrahydrofolate ligase and the knockout of GHMT2r and GLYCL was applied (*i*JN1463_LBB05). In addition, the growth-coupling of 3HK production from L-tryptophan using glucose as the sole carbon source was confirmed by adding the 3HK synthesis pathway to *i*JN1463_LBB05, resulting in *i*JN1463_LBB07.

The ability to produce xanthommatin was calculated using model simulations. Therefore, a lumped reaction representing dimerase and oxidoreductase activity was added to *i*JN1463_LBB07 (*i*JN1463_xtm), which converts 3HK to xanthommatin at the stoichiometric expense of oxygen and the release of ammonia and water. An alternative xanthommatin reaction scheme was additionally tested using NADP^+^ as the electron acceptor instead of oxygen.

All scripts and metabolic models used to run GCS and conduct model analyses are provided in the Supplementary Appendix. Throughout this study, the COBRApy metabolic modeling package toolbox (version 0.20.0)^59^ in combination with the Gurobi Optimizer suite (version 9.0.0)^60^ were used for model handling and simulations. The GCS workflow was run on the high-performance computing cluster of the Technical University of Denmark (DTU) with a maximum configuration of 48 GB of RAM and 24 Intel Xeon 2660v3 processors (2.60 GHz). Data processing and model analysis routines were implemented and run in Python 3.9 on a Windows 10 machine with 32 GB of RAM and an AMD Ryzen 5900x processor (12 cores at 3.7 GHz).

### Growth assays in microplates

A pre-culture was prepared by inoculating a single colony into 3 mL LB medium in a 14 mL culture tube. After growth for 16 – 24 hours, a 100 μL-aliquot of culture was passaged to a new 14 mL culture tube containing DBM medium supplemented with 10 mM formate. After growth overnight, cells were harvested by centrifugation (13000g for 2 min), washed twice in DBM, and diluted into DBM medium to OD_600_ = 0.1 - 0.2. The cell suspension (50 *µ*L per well) was distributed into a 96-well clear plate that contained DBM medium (50 *µ*L per well), giving an initial OD_600_ = 0.05-0.1. DBM was supplemented with glycine, formate, L-tryptophan, and antibiotics according to the strain requirements and experimental conditions. Following setup, plates were covered with Breathe-Easy sealing membrane. Cell density (OD_600_) was measured in a Tecan Spark plate reader using a kinetic cycle of 12 shaking steps, which alternated between linear and orbital (1 mm amplitude), each 60 s long. Experiments were performed in two or three biological replicates. The growth curves shown represent the average of replicates.

### Bioproduction assays in shake flasks

For each replicate, an initial pre-culture was prepared by inoculating a single colony into 3 mL LB (+Sm) in a 15 mL culture tube. After growth for 16 – 24 h, a 100 μL-aliquot of cells was diluted into 3 mL DBM (+Sm), supplemented with 10 mM formate or 5 mM L-tryptophan. Following overnight growth, a 2% inoculum was transferred into a 125 mL-shake flask containing 25 mL DBM (+Sm). DBM medium contained glycine and L-tryptophan supplements as defined by the experiment. Cultures were incubated at 30°C, 225 r.p.m. for 3 days. At defined timepoints, a 0.5 mL aliquot was withdrawn from the culture and stored at -80°C for subsequent metabolite analysis. At each timepoint, optical density was measured using an Agilent Cary 60 UV-Vis spectrophotometer. Two biological replicates were tested per strain.

### Metabolite analysis by HPLC

Frozen culture samples were thawed and centrifuged (16000g for 2 min). The supernatant was collected, filtered through a 0.22 μm filer, and analyzed using an Agilent LC/MSD iQ instrument consisting of an Agilent 1290 Infinity Series HPLC system, an automated liquid sampler, a diode array detector, and a temperature-controlled column compartment. The mobile phases were water (solvent A) and MeCN (solvent B), both containing 0.1% (v/v) formic acid (FA). Next, 10 *µ*L-aliquots of filtered samples were injected onto a Phenomenex Kinetex C18 100-Å column (2.6 *μ*m, 4.6 mm × 100 mm), equipped with a guard column, operating at 0.5 ml min^−1^. Elution was performed with an isocratic step of 10% B for 3 min, followed by gradient steps of 10–30% B over 11 minutes and 30–100% B over 4 minutes. OpenLabs CDS software was used to operate the instrument and to analyze data. Sample concentrations were quantified by comparing to calibration curves of known concentrations.

### Xanthommatin identification by HPLC-coupled HR-MS

Filtered culture supernatant was treated with 1% ascorbic acid and incubated at 4°C overnight. The resulting precipitant was collected by centrifugation (16000g for 5 min), re-suspended in an equal volume of MeOH (+1% (v/v) HCl), and filtered through a 0.22 μm filer. Samples were analyzed using an Agilent 6530 quadrupole time-of-flight (Qtof) MS instrument, consisting of an automated liquid sampler, a 1260 Infinity Series HPLC system, a diode array detector, an ESI source and the 6530 Series Qtof. The mobile phases were water (solvent A) and MeCN (solvent B), both containing 0.1% formic acid (FA). Samples were injected onto a Phenomenex Kinetex C18 100-Å column (2.6 μm, 4.6 mm × 100 mm) equipped with a guard column operating at 0.5 ml min^−1^. Elution was performed with an isocratic step of 10% B for 3 min, followed by gradient steps of 10–30% B over 11 minutes and 30–100% B over 4 minutes. MassHunter software was used to operate the instrument and to analyze data.

### Adaptive laboratory evolution

ALE experiments were performed on a previously described automated platform^61^ following a previously described selection pressure protocol to adapt organisms to growth on non-native substrates.^50^ In brief, strains were inoculated into DBM containing 20 mM glucose and growth-enabling supplements of either 5 mM L-tryptophan or 5 mM glycine with 10 mM formate. Culture growth was periodically monitored by measuring optical density at a 600 nm wavelength using a Tecan Sunrise microplate absorbance reader. Once cultures had reached the stationary growth phase for a period of at least 1 day, 300 *µ*L was propagated into each of two batches. The first contained 0.5 mM less of each of the growth-enabling supplements than the previous batch, and the second “test” batch contained no growth-supplements except for 5 mM of formate in the glycine auxotroph removal ALE. If the test flask reached an OD_600_ of at least 0.05 and either hit stationary phase or 4 consecutive days of culturing, it was considered to be growing and 300 *µ*L was propagated to a new batch. After at least two consecutive batches showed growth, the adaptation was considered successful. If an OD_600_ of 0.05 was not reached within 4 days, the test batch was terminated, and a new one was started the next time the growth-enabling supplemented line reached stationary phase. All ALE replicate lineages were started by inoculation from independent colonies isolated on agar plates with the same media composition as the ALE experiment. Replicate lineages were grown in 17 mL batches maintained at 30°C ± 0.3°C. Batches were well aerated by vigorous stirring at 1200 rpm. Endpoint populations were streaked out from frozen glycerol stocks onto LB agar to obtain isolated clones. Clones were chosen, at random, and tested for the desired phenotype in microplate-based growth assays. Select clones were subjected to whole-genome sequencing.

### Whole-genome sequencing and mutation analysis

Strains were cultured overnight in LB containing appropriate supplements. 2 mL of culture was harvested by centrifugation for DNA extraction. Cell pellets were extracted immediately or stored at -80°C. Methods for DNA extraction, library preparation, and sequencing are detailed in Supplementary Table 4. Sequencing services were performed by UC San Diego Institute for Genomic Medicine or Plasmidsaurus. The sequencing results were processed with the in-house pipeline that incorporates the BreSeq^62^ pipeline to identify mutations. The *Pseudomonas putida* KT2440 genome sequence (GenBank Accession AE01545) was used as a reference sequence.

### Linear amplicon sequencing and variant analysis

<1 *µ*L glycerol stock of evolved population A8-F3-I0 was PCR amplified using Q5 Master Mix and primers metK_g-check_1 and metK_g-check_4. The resulting PCR product was purified and subjected to Premium PCR Sequencing (Primordium Labs). The sequencing data was upload to the Galaxy web platform and the public server at usegalaxy.org was used to analyze the data.^63^ Raw reads in fastq format were aligned to the reference gene using BowTie2 with the analysis mode set to default settings.^64^ Bam files from Bowtie2 were then used to generate pileup formatted alignments using Samtools mpileup.^65^ Genetic variants (SNPs, indels) were analyzed using VarScan mpileup using the single nucleotide variation analysis with the following settings: Minimum coverage=1, Minimum supporting reads=3, Minimum base quality=15, Minimum variant allele frequency =0.01, Minimum homozygous variant allele frequency =0.75, Default p-value threshold for callings variants =0.99. VCF files from VarScan were combined into a single file which was used to determine the frequency and position of mutational variants.^66^

### Isolation of DC-Xa for NMR analysis

Biomass from an LB agar plate was used to inoculate a seed culture in LB-Sm which was then grown for 16 – 24 h and used to inoculate 2.8 L-fernbach shake flasks containing 500 mL DBM at 2% dilution. After growth at 30°C, 225 r.p.m for 72 h, cultures were centrifuged (for 20 min). Supernatants were collected and passed through a 0.22 μm vacuum filter system to remove cells and debris. After addition of ascorbic acid at a final concentration of 1% (w/v), the supernatant was incubated overnight at 4°C. The next day, the precipitate was collected by centrifugation, washed twice with ddH_2_O, and dried by lyophilization. Lyophilized material was resuspended in MeOH with 1% (v/v) HCl before being subjected to semi-preparative RP-HPLC using an Agilent 1260 Infinity system equipped with a Phenomenex Luna C18(2) column (5 *µ*m, 100 Å, 250 × 10 mm). The separation was carried out using a gradient of 20–40% acetonitrile (CH_3_CN) with 0.1% (w/v) TFA over 20 minutes. DC-Xa was collected at a retention time (tR) of 9.4 min. The collected fraction was further purified on semi-preparative RP-HPLC on an analytical Eclipse XDB-C18 (5 *µ*m, 4.6 × 150mm) using a gradient of 10-35% CH_3_CN + 0.1% (w/v) TFA (0–20 min), yielding a DC-Xa peak at t_R_ 8.2. This fraction was subsequently repurified on the same Eclipse XDB-C18 column using an isocratic 10% CH_3_CN + 0.1% TFA method, collecting the final purified DC-Xa at tR 10.8 minutes. Approximately 2 mg of purified DC-Xa was dissolved in 600 *µ*L d4-DMSO (+4% d1-TFA) and transferred into a 5 mm tube. NMR spectra were acquired at UCSD Skaggs School of Pharmacy NMR facilities on a Bruker Avance III (600 MHz) NMR spectrometer with a 1.7 mm ^1^H[_13_C/_15_N] microcryoprobe (Billerica, MA).

### Fed-batch fermentation in bioreactor

Biomass of ePUMA-Xa(WIn) (clone A18.F4.I5) from an agar plate was used to inoculate a pre-culture in LB-Sm that was grown overnight. This culture was used to inoculate a second preculture of 10 mL DBM medium with 20 mM glucose, 10 mM sodium formate and Sm incubated for 24 h at 200 r.p.m. The final preculture was done in a 250 mL Erlenmeyer flask containing 30 mL of DBM with 20 mM Glucose and Sm, which was incubated at 30°C and 200 rpm. After 16 h, the culture was centrifuged and resuspended in 5 mL DBM such that the initial starting OD_600_ was 0.01 in 160 mL of total culture in a 250 mL AMBR reactor; 20% (v/v) orthophosphoric and 1 M NaOH were used as titrants to regulate the pH to 7. The stirrer was set to 500 rpm., increasing to 1800 r.p.m. at the end of the fermentation. Exponential feeding was chosen on 0.15003 mL/h based on the specific growth rate of 0.1667 h^-1^, glucose concentration of 400 g/L in the feeding bottle and the yield 0.2625 g_CDW_/g glucose. Fed-batch was triggered by a sudden increase in the dissolved oxygen to a range of 30-40%. Fermentation lasted 50 h, when the maximum working volume was reached, during that time samples of 1 mL were automatically taken every 6 h. Sample processing consisted of measuring OD_600_ via spectrophotometer, followed by centrifugation to obtain supernatant for further HPLC analysis of carbon source and products. The final volume was centrifuged at 4000 r.p.m. for 20 min, then supernatant was filtered through a 0.2 *µ*m filter to start the purification process with ascorbic acid described above.

### Chemical synthesis of Xanthommatin

Synthetic xanthommatin (Xa) was synthesized from 3-hydroxy-DL-kynurenine (3-HK, Sigma-Aldrich) using potassium ferricyanide (Thermo Scientific) according to previous protocols.^10,52,67^ After 90 min, the product was precipitated with 90 μL of 1 M hydrochloric acid (Sigma-Aldrich), which was added dropwise to the mixture to crash out the product from the reaction solution. The resulting precipitate was washed with deionized (DI) water three times after centrifugation at 5000g for 5 minutes.

### Analytical characterization of xanthommatin

UV-vis absorbance spectra (Thermo Scientific, Evolution 220 UV-visible spectrophotometer) were recorded by dissolving 0.15 mg/mL of each sample in dimethyl sulfoxide (DMSO, Fisher). Redox-dependent color change was monitored in both the synthetic Xa and biosynthetic Xa solutions containing 0.25 mg of each material in 200 μL of 0.1 M 2-(*N*-morpholino)ethanesulfonic acid (MES) buffer. Here, 10 μL of an oxidant (1 M sodium nitrite in DI water, Alfa Aesar) was added to oxidize the product. Conversely, the addition of reductant (1 M ascorbic acid in DI water, Sigma-Aldrich) was added to reduce the product. Cyclic voltammetry curves were obtained using Gamry Interface 1000 B in a three-electrode configuration containing a glassy carbon working electrode, a platinum wire counter electrode, and silver-silver chloride (Ag/AgCl with saturated KCl) reference electrode. The supporting electrolyte consisted of 3 mL of DMSO with 0.1 M lithium triflate (Sigma-Aldrich), in which 0.15 mg/mL of each sample was dissolved. The voltammograms were recorded over five cycles, scanning from -0.2 to 0.6 V at a rate of 50 mV/s, with the third cycle represented.

Based on the obtained absorption spectra, the unique electronic transition and energy bandgaps for each compound can be extrapolated using the Tauc plot. To construct Tauc plots for determining the energy bandgaps of prepared Xa, the direct energy bandgap of ((αhv)^2^ vs E_g_) was compared. E_g_ was extrapolated from the linear segments in the plot. The initial transition at lower energy represents the optical bandgap, which marks the initial formation of an electron-hole pair. The subsequent transition is referred to as the fundamental energy gap, which denotes the energy difference between the valence band (or highest occupied molecular orbitals, HOMO) and the conduction band (or lowest unoccupied molecular orbitals, LUMO).^68^ Analysis of the optical and fundamental energy bandgaps revealed crude Xa has optical and fundamental bandgaps of 2.49 eV and 3.71 eV, respectively, while the biosynthesized Xa exhibits optical and fundamental bandgaps of 2.43 eV and 3.78 eV, respectively. While slightly different, the bandgaps of the two samples are comparable and consistent with our previous reports on synthetic Xa in materials.^8^

### Data analysis

Data handling and calculations were performed in Microsoft Excel and OriginPro (OriginLab). Figures and illustrations were created in OriginPro and Adobe Illustrator. Geneious Prime was used to design and manage DNA constructs and to analyze sequencing results.

