## Supplementary Information for "Growth-coupled microbial biosynthesis of the animal pigment xanthommatin"

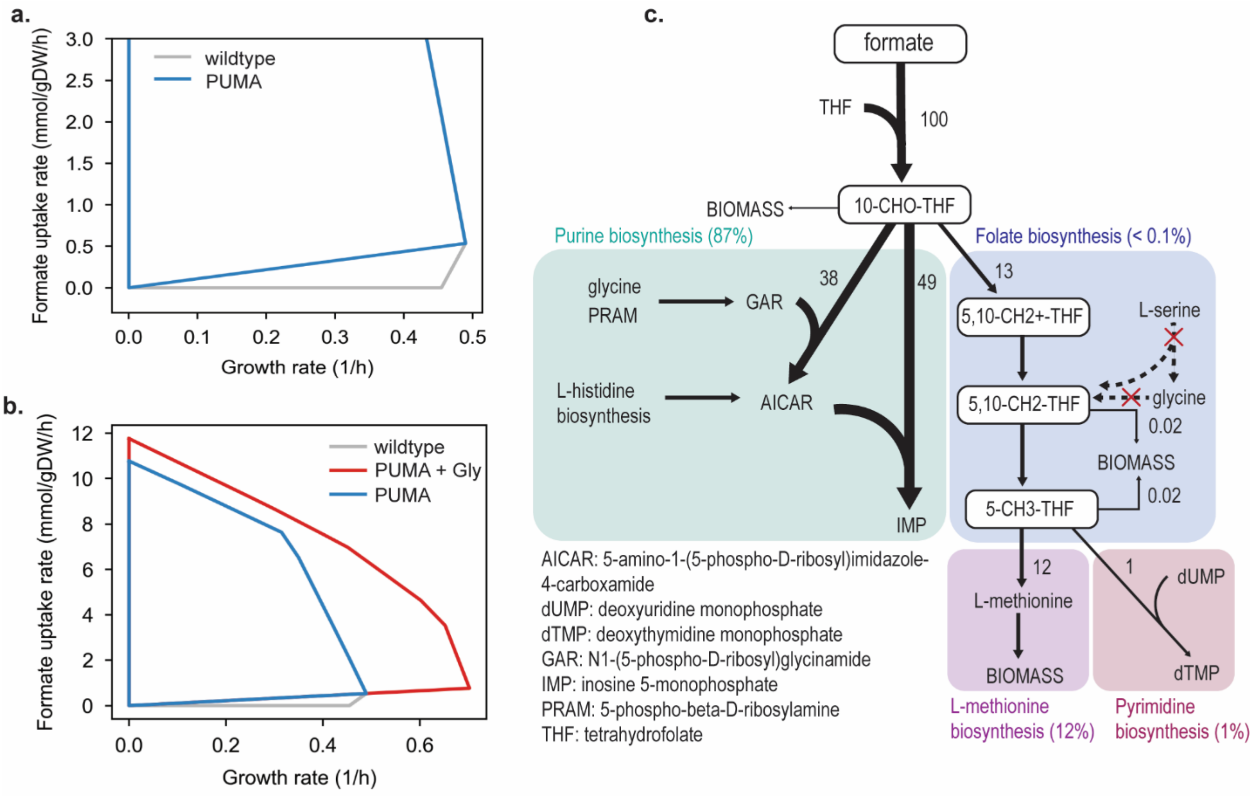


**Supplementary Figure. 1:** Metabolic model-based analysis of C1 growth-coupled biosynthetic (GrowBio) strategy; **a.** 2D flux spaces spanned by growth rate and formate uptake rate for a *P. putida* wildtype strain (*i*JN1463) and its derivative PUMA including the heterologous formate assimilation pathway and the knockout of the glycine cleavage system (*gcvTHP*) and serine hydroxymethyltransferase (*glyA*); **b.** Flux space of the PUMA strain with a glycine supplementation is additionally shown; **c.** Metabolic destinations of the derivatives from the formate assimilation in the PUMA metabolic model and a corresponding flux distribution are shown. Fluxes are given relative to an arbitrary formate uptake rate of 100. A glucose minimal medium was applied under aerobic conditions for all simulations.

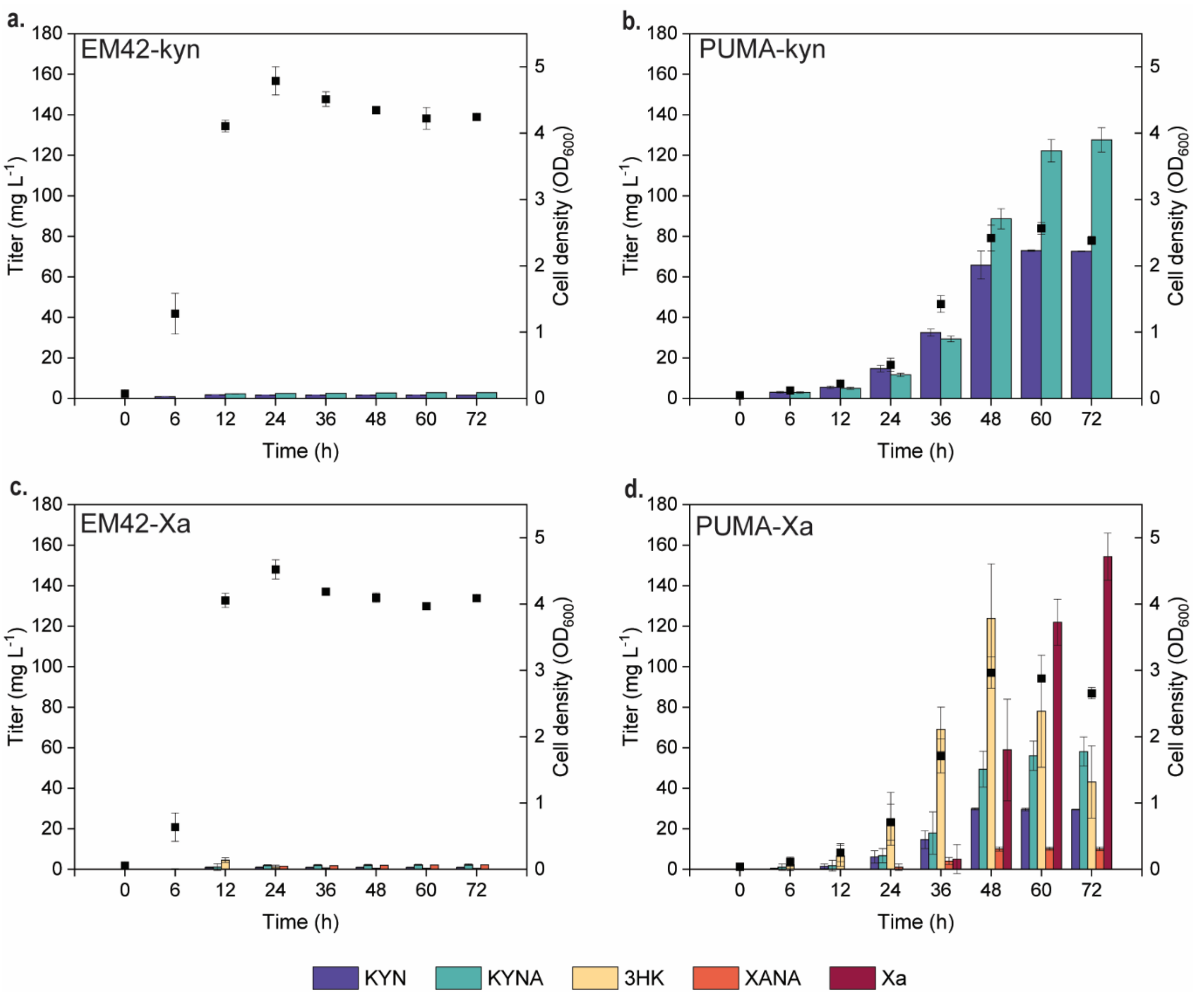


**Supplementary Figure 2.** Metabolite production of engineered strains. Strains were cultivated in shake flasks filled with 20% (v/v) DBM medium supplemented with 20 mM glucose, 5 mM glycine and 5 mM L-tryptophan. Averages and standard deviations are presented for two biological replicates.

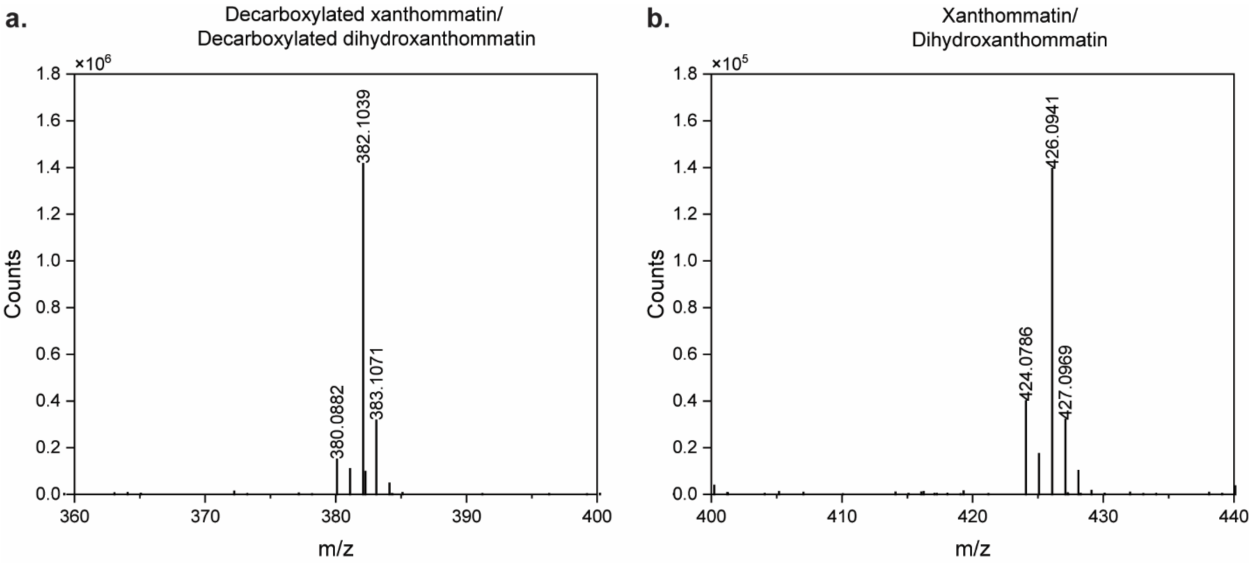


**Supplementary Figure. 3.** Identification of xanthommatin compounds by high resolution mass spectrometry.

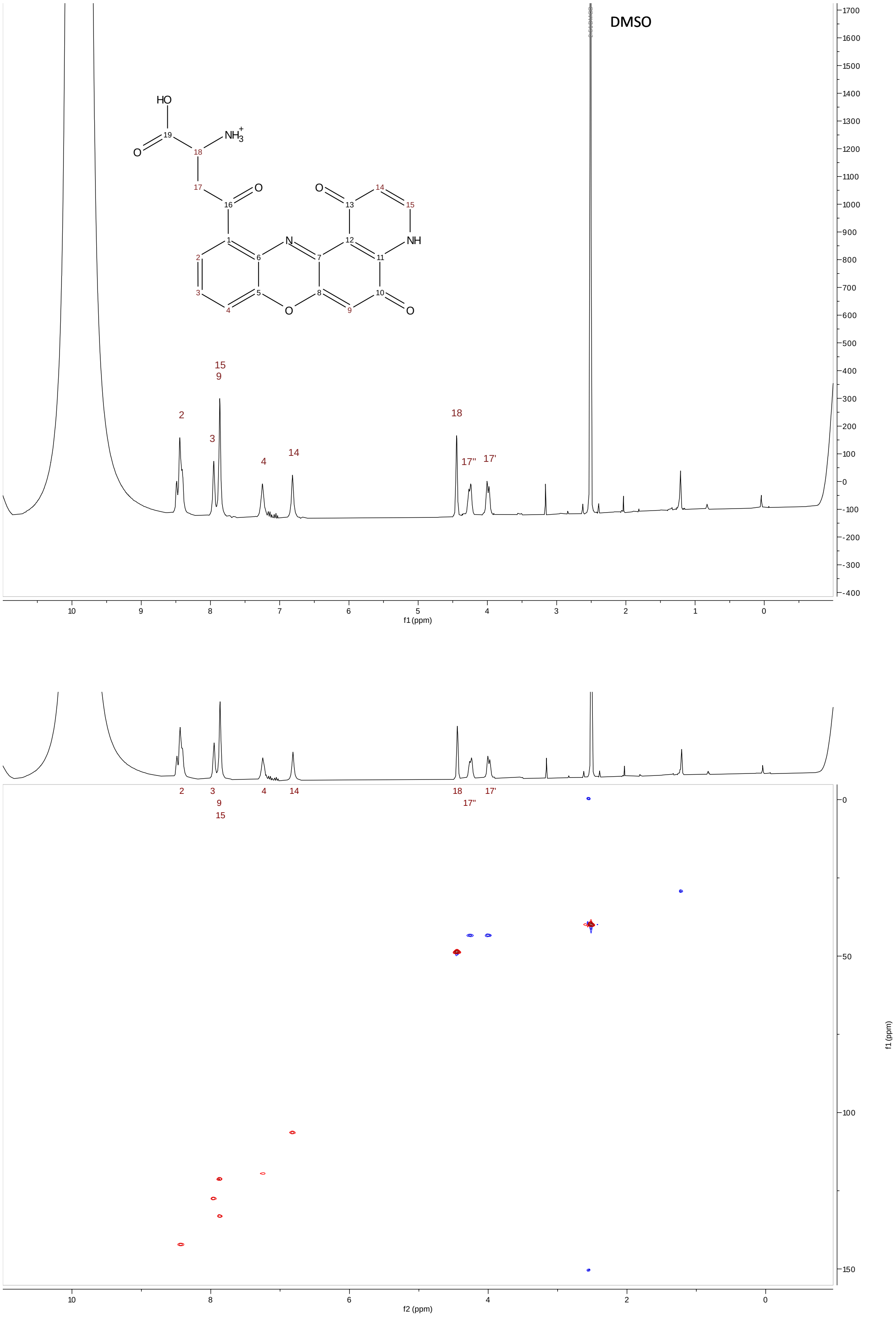


**Supplementary Figure 4**. NMR spectra of decarboxylated xanthommatin in d6-DMSO (+11% d1-TFA). Shown are: $^1$H (top) and HSQC (bottom) spectra.

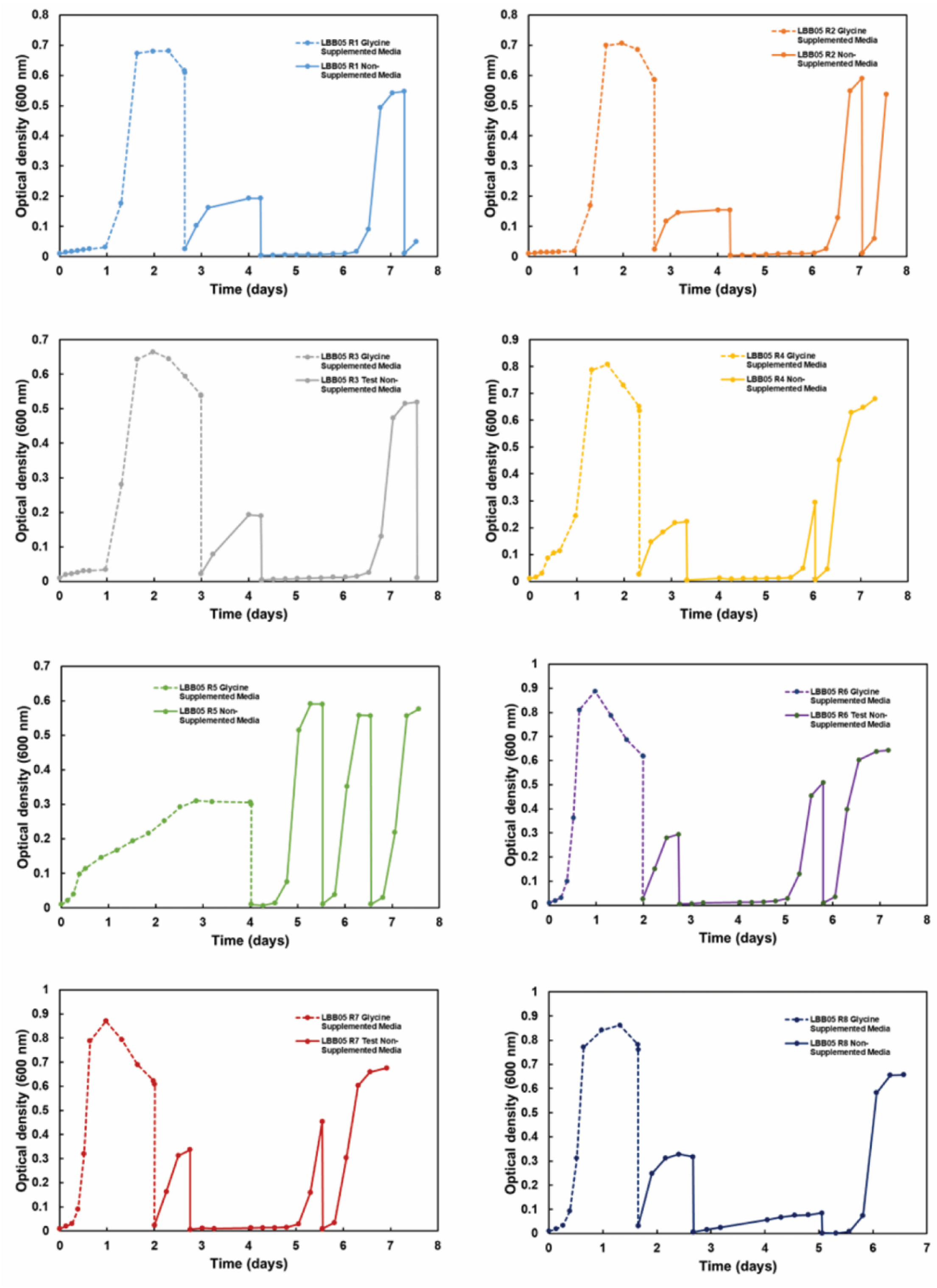


**Supplementary Figure 5.** Growth curves for the glycine auxotroph removal ALE experiment. All replicates showed consistent growth on the base media after a single batch of growth in glycine-supplemented media. Clonal isolates for all replicates were isolated from the last batch of growth in the non-supplemented media.

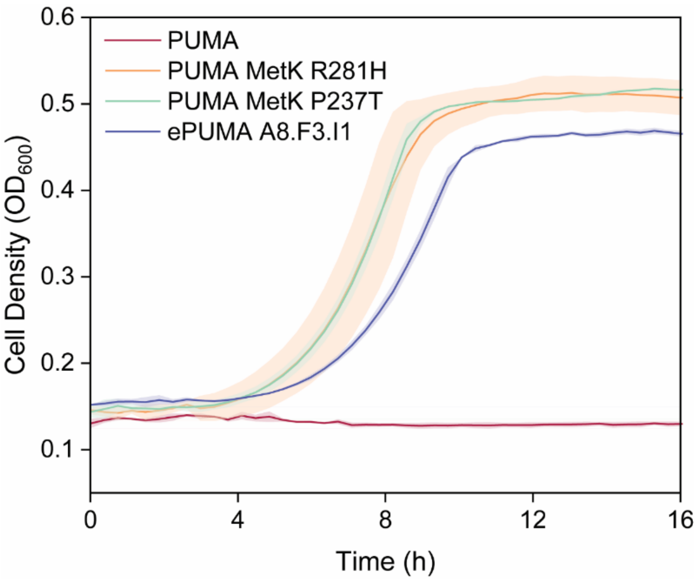


**Supplementary Figure. 6.** A growth plot of the PUMA strain and its derivatives in DBM medium containing glucose (20 mM) and formate (5 mM) without supplemental glycine. PUMA is not able to grow in the absence of glycine whereas ‘re-engineered’ PUMA mutants show growth comparable to an ePUMA clonal isolate. Average values and standard deviation of two biological replicates are presented.

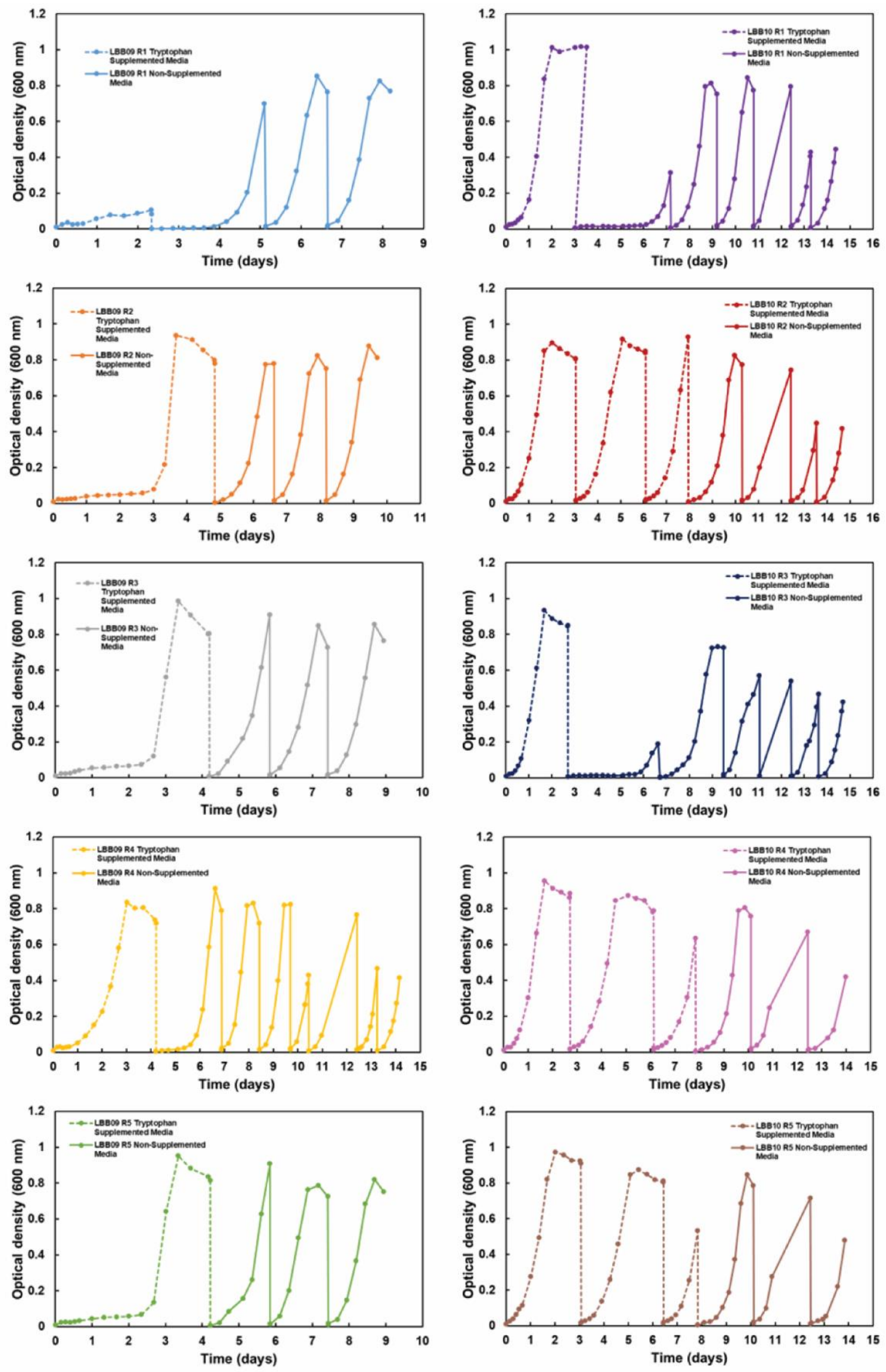


**Supplementary Figure. 7.** Growth curves for the tryptophan auxotroph removal ALE experiment. All replicates showed consistent growth on the base media after no more than three batches of growth in tryptophan-supplemented media. Clonal isolates for all replicates were isolated from the last batch of growth in the non-supplemented media.

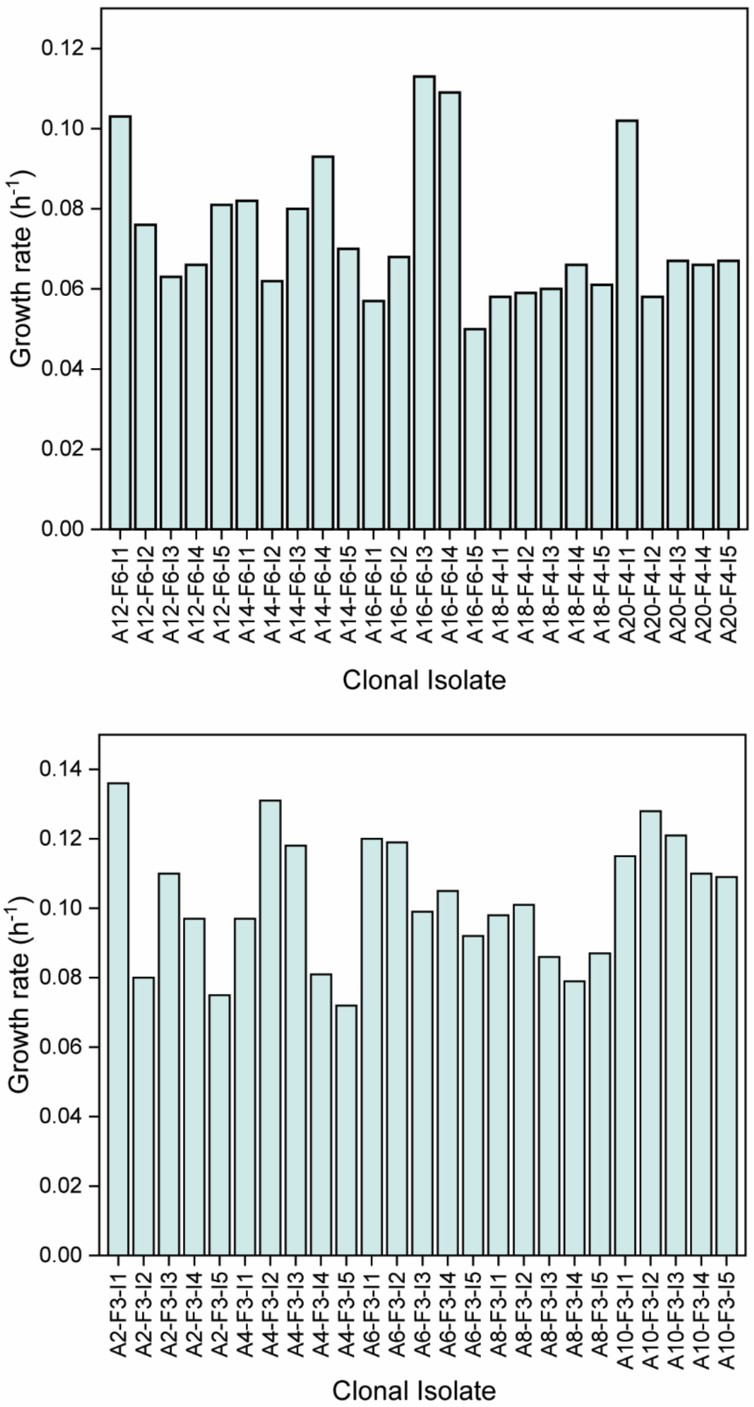


**Supplementary Fig. 8.** Growth rate characterization of single clones from evolved populations (5 clones per lineage) obtained via adaptive laboratory evolution of ePUMA-kyn (bottom) and ePUMA-Xa (top). Strains were cultivated in 96-well microtiter plates in DBM medium supplemented with 20 mM glucose. All 50 clonal isolates screened showed growth in the absence of supplemental L-tryptophan.

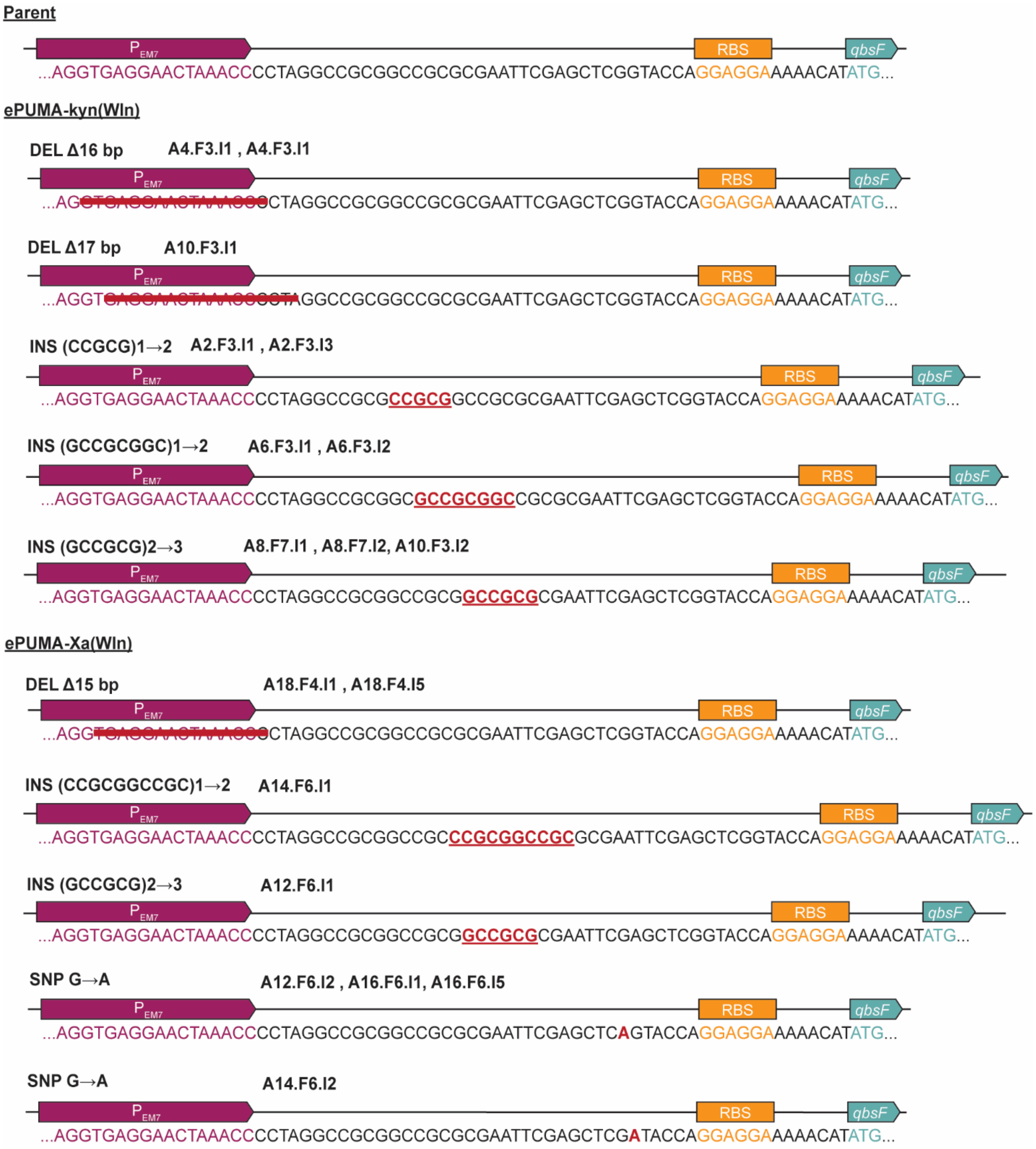


**Supplementary Fig. 9.** Sequences of promoters in evolved pS4413-*qbsFH* and pS4413-*qbsFGH* plasmids. Whole-genome sequencing of clonal isolates from endpoint populations of ePUMA-kyn and ePUMA-Xa, weaned from L-tryptophan supplementation, revealed mutations in the promoter region controlling expression of *qbs* biosynthetic genes.

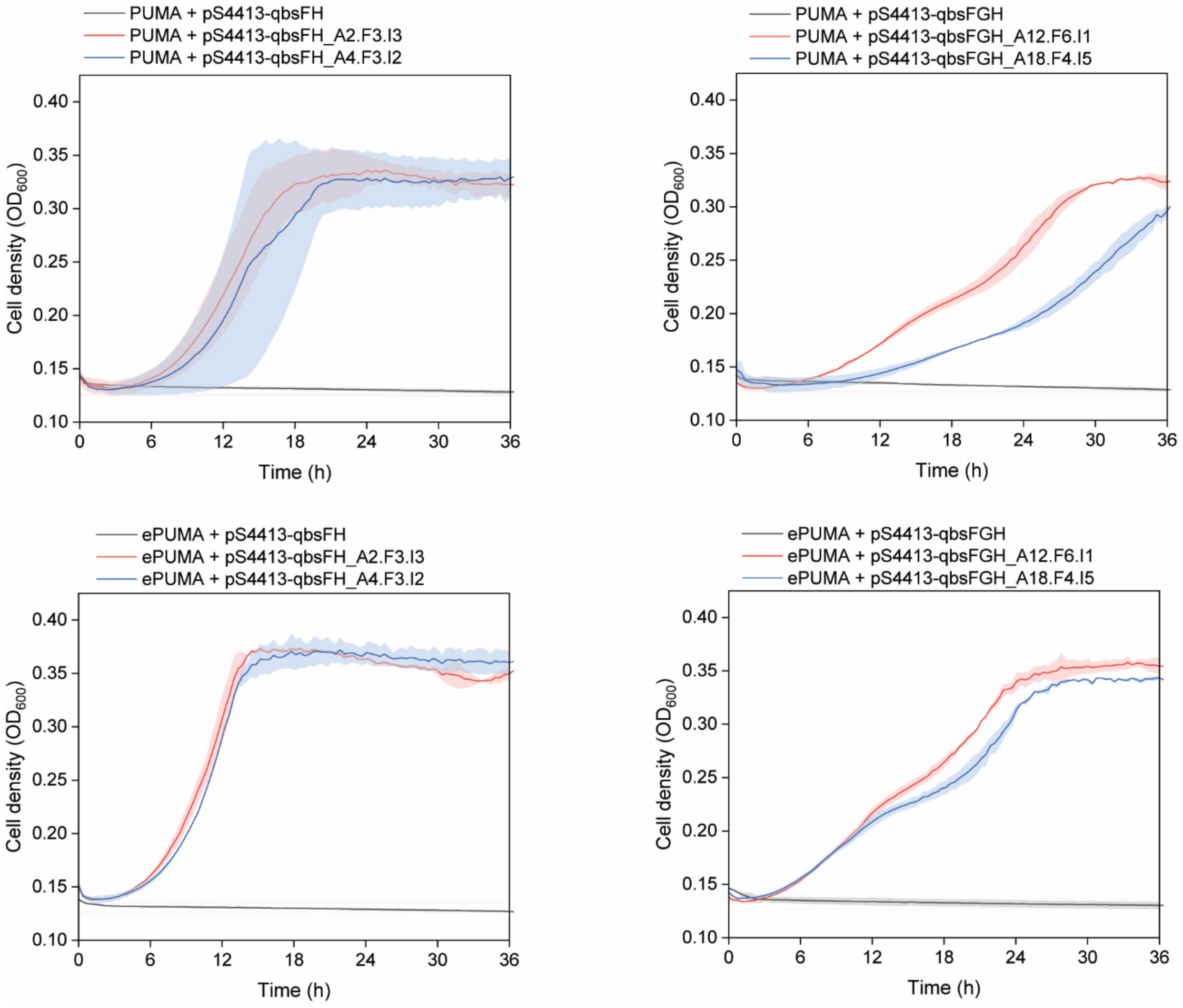


**Supplementary Figure 10.** Re-engineering evolved pathway plasmids pS4413-*qbsFH* and pS4413-*qbsFGH* into PUMA and ePUMA. Mutant plasmids were isolated from single clones (ALE IDs indicated) of ePUMA-kyn(WIn) and ePUMA-Xa(WIn) clones and transformed into the PUMA and ePUMA strains. The PUMA strains harboring the mutant plasmids were able to grow in DBM medium containing glucose (20 mM) and supplemental glycine (5 mM) in the absence of L-tryptophan whereas the PUMA strain carrying the original pS4413-qbsFH or pS4413-qbsFGH plasmid did not. Similary, ePUMA only grew in DBM medium containing glucose as the sole carbon source when carrying mutant pS4413-*qbsFH* and pS4413-*qbsFGH* plasmids, but the originals.

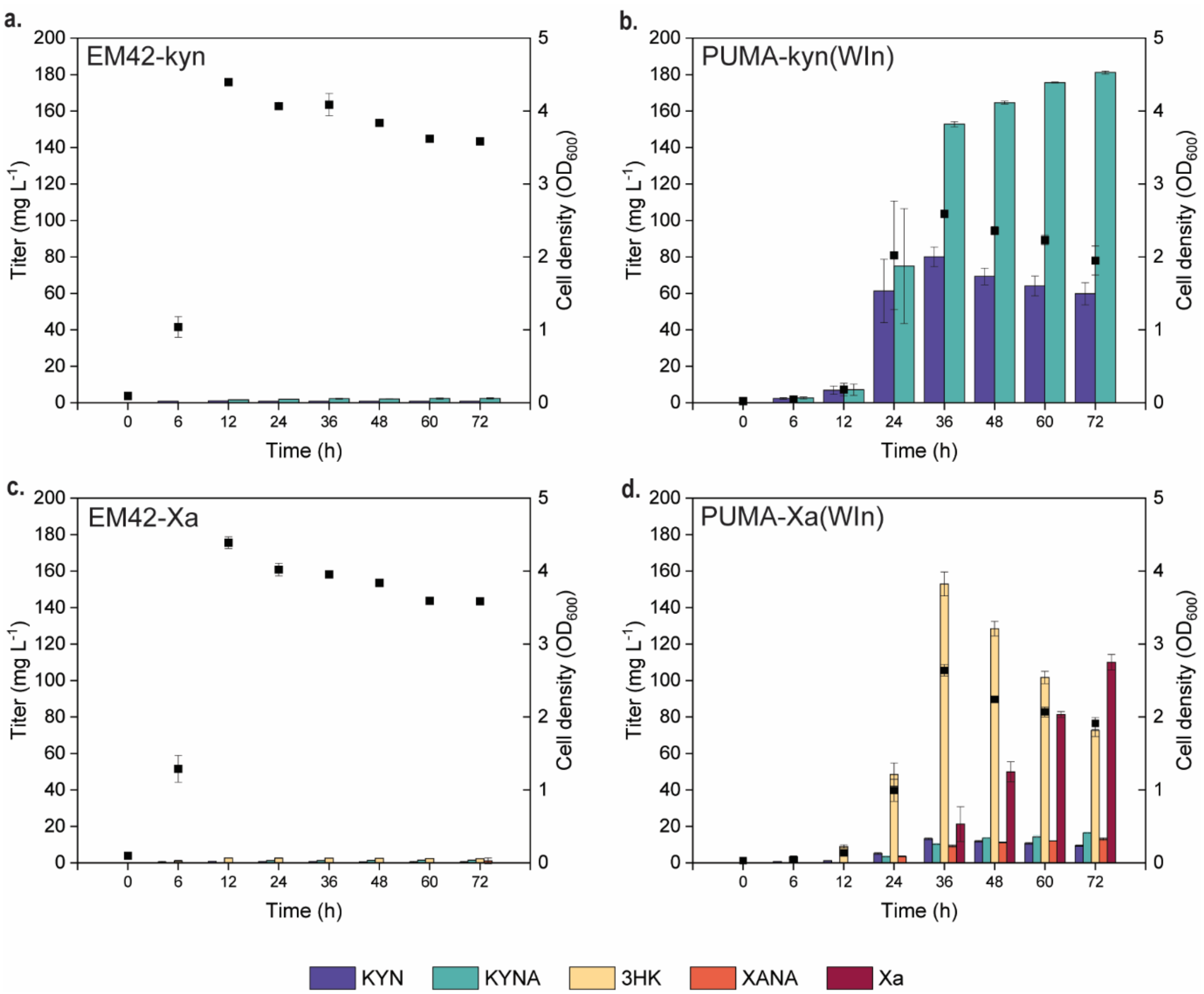


**Supplementary Figure 11.** Metabolite production of engineered strains from glucose as the sole carbon source. Strains were cultivated in shake flasks filled with 20% (v/v) DBM medium supplemented with 20 mM glucose. Averages and standard deviations are presented for two biological replicates.

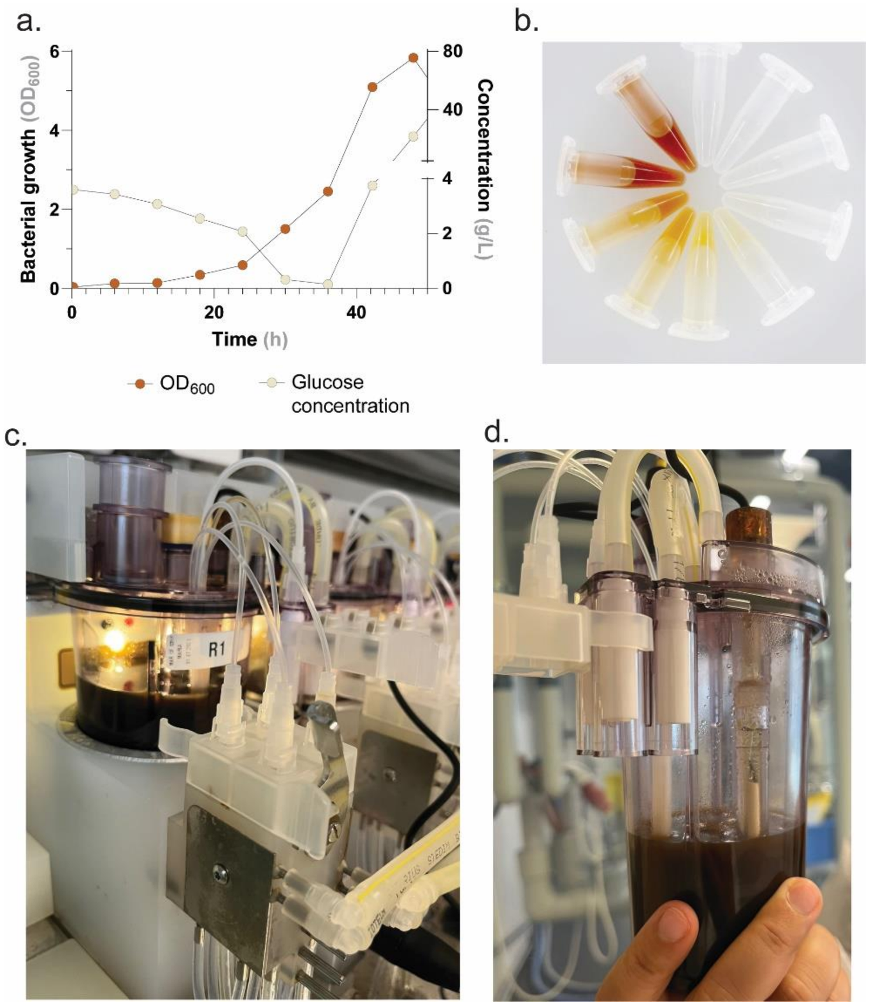


**Supplementary Figure 12.** Fed-batch fermentation of PUMA-Xa(WIn). a. A growth plot showing glucose concentration over the course of the fermentation. b. Samples taken every 6 h from the fermentation broth show a visual increase in xanthommatin production as indicated by the yellow to brown color change. c. and d. Fermentation was performed in a 250 mL Ambr microbioreactor. The final culture broth was colored deep burgundy.

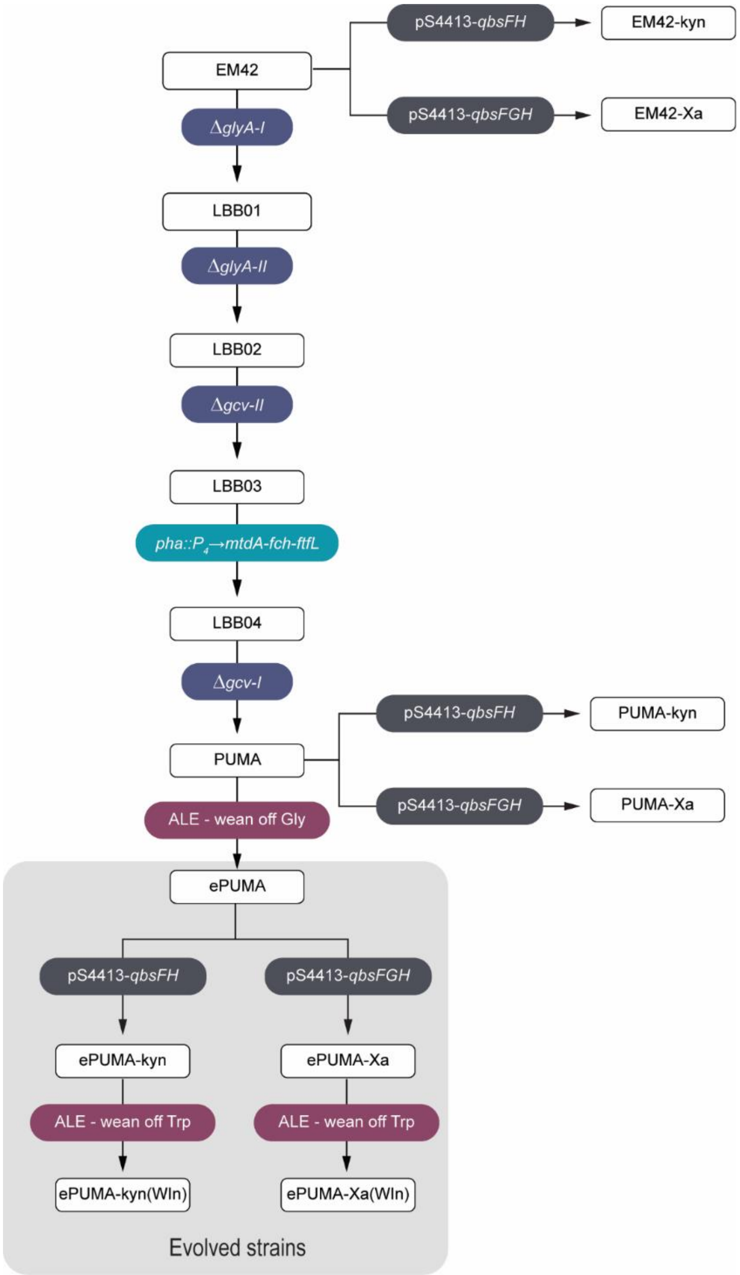


**Supplementary Figure 13.** Genealogy of strains created in this manuscript. A *P. putida* methylenetetrahydrofolate auxotroph was generated from P. putida EM42 via five genomic modifications, which included deletion of *glyA-I*, *glyA-II*, and *gcv-II*, insertion of a formate assimilation module (P4→*mtdA-fch-ftfL*) and deletion of *gcv-I*, in that order. The resulting strain named PUMA was subjected to adaptive laboratory evolution (ALE) to remove its dependence on exogenous glycine for growth in minimal medium. One of the evolved populations of the resulting glycine-independent strain, named ePUMA, was transformed with the pathway plasmids pS4413-*qbsFH* or pS4413-*qbsFGH*, to generate ePUMA-kyn and ePUMA-Xa, respectively. Single transformants were subsequently subjected to ALE in parallel to remove the dependence on exogenous L-tryptophan for growth in minimal medium. The resulting strains ePUMA-kyn(WIn) and ePUMA-Xa(WIn) grow in minimal medium containing glucose as the sole carbon source.

**Supplementary Table 1.** Strains used in this study.

| Strain | Genotype/Relevant characteristics | Reference or source |
|---|---|---|
| *Escherichia coli* | | |
| DH5α | Cloning host; F⁻ λ⁻ *endA1 glnX44*(AS) *thiE1 recA1 relA1 spoT1 gyrA96*($Nal^R$) *rfbC1 deoR nupG* Φ80(*lacZ*ΔM15) Δ(*argF-lac*)*U169 hsdR17*($r_k^-$ $m_K^+$) | Hanahan and Meselson (1983) |
| DH5α λpir | Cloning host; F⁻ λ⁻ *endA1 glnX44*(AS) *thiE1 recA1 relA1 spoT1 gyrA96*($Nal^R$) *rfbC1 deoR nupG* Φ80(*lacZ*ΔM15) Δ(*argF-lac*)*U169 hsdR17*($r_k^-$ $m_K^+$), λ pir lysogen | Platt et al. (2000) |
| *Pseudomonas putida* | | |
| KT2440 | Wild-type strain, derived from *P. putida* mt-2 (Williams and Murray, 1974) cured of the TOL plasmid pWW0 | Bagdasarian et al. (1981) |
| EM42 | Reduced-genome derivative of P. putida KT2440; ΔPP_4329-PP_4397 (flagellar operon) ΔPP_3849-PP_3920 (prophage I) ΔPP_3026-PP_3066 (prophage II) ΔPP_2266-PP_2297 (prophage III) ΔPP_1532-PP_1586 (prophage IV) ΔTn7 ΔendA-1 ΔendA-2 ΔhsdRMS ΔTn4652 | Martínez-García et al. (2014b) |
| LBB01 | EM42 Δ*glyA-I* (PP_0322) | This study |
| LBB02 | LBB01 Δ*glyA-II* (PP_0671) | This study |
| LBB03 | LBB02 *gcv-II* (PP_5192-PP_5194) | This study |
| LBB04 | LBB03 pha::P4→mtdA-fch-ftfL ΔPP_5002-PP_5008:: P4→mtdA-fch-ftfL from *M. extorquens* | This study |
| PUMA (LBB05) | LBB04 Δgcv-I (PP_0986-PP_0989) | This study |
| PUMA-kyn (LBB06) | LBB05 pS4413-*qbsFH* | This study |
| PUMA-Xa (LBB07) | LBB05 pS4413-*qbsFGH* | This study |
| ePUMA | Glycine-independent PUMA derivative | This study |
| ePUMA-kyn | ePUMA pS4413-*qbsFH* | This study |
| ePUMA-Xa | ePUMA pS4413-qbsFGH | This study |
| EM42-kyn | EM42 pS4413-*qbsFH* | This study |
| EM42-Xa | EM42 pS4413-*qbsFGH* | This study |
| ePUMA-kyn(WIn) | Tryptophan independent ePUMA-kyn derivative | This study |
| ePUMA-Xa(WIn) | Tryptophan independent ePUMA-kyn derivative | This study |
| PUMA MetK R281H | PUMA *metK* R281H (CGC→CAC) | This study |
| PUMA MetK P237T | PUMA *metK* P237T ((CCG→ACG) | This study |
| *Pseudomonas* sp. | | |
| DTU12.1 | Quinolobactin producer; source of *qbsFGH* genes | Sazinas et al. (2019) |

**Supplementary Table 2.** Plasmids used in this study.

| Plasmid | Relevant characteristics | Reference or source |
|---|---|---|
| pGNW2·ΔglyA-I | Derivative of vector pGNW2 carrying HRs to delete *glyA-I* (PP_0322); Km$^R$ | Turlin et al. (2022) |
| pGNW2·ΔglyA-II | Derivative of vector pGNW2 carrying HRs to delete *glyA-II* (PP_0671);Km | Turlin et al. (2022) |
| pGNW2·Δgcv-I | Derivative of vector pGNW2 carrying HRs to delete *gcv-I*; Km$^R$ | Turlin et al. (2022) |
| pGNW2·Δgcv-II | Derivative of vector pGNW2 carrying HRs to delete *gcv-II*; Km$^R$ | Turlin et al. (2022) |
| pGNW2·pha::M1 | Suicide vector for integration of a P4→*mtdA-fch-ftfL* module from M. extorquens into the native *pha* (PP_5002-PP_5008) locus of *P. putida* | Turlin et al. (2022) |
| pQURE6·H | Conditionally-replicating vector carrying XylS/Pm→I-SceI and P14g(BCD2)→mRFP; GmR | Volke et al. (2020b) |
| pS448·CsR | Derivative of vector pSEVA448 used for CRISPR-Cas9 counterselection; xylS (cured of BsaI restriction sites), Pm→cas9, $P_{EM7}$→sgRNA; Sm$^R$ | Wirth et al. (2019) |
| pS448·CsR_glyA-II_2 | Derivative of vector pS448·CsR carrying $P_{EM7}$→*glyA-II*-targeting sgRNA; Sm$^R$ | This study |
| pS448·CsR_gcv-I_2 | Derivative of vector pS448·CsR carrying $P_{EM7}$→*gcv-I*-targeting sgRNA; Sm$^R$ | This study |
| pSEVA4413 | Expression vector; $P_{EM7}$→MCS (for constitutive gene expression), *oriV*(pRO1600/ColE1); Str$^R$ | Silva-Rocha et al. (2013) |
| pS4413-qbsFH | Derivative of vector pSEVA4413 carrying the *qbsF* and *qbsH* genes from *Pseudomonas* sp. DTU, Sm$^R$ | This study |
| pS4413-qbsFGH | Derivative of vector pSEVA4413 bearing the *qbsFGH* casette from *Pseudomonas* sp. DTU, Sm$^R$ | This study |
| pS4413-qbsFH_A2.F3.I3 | Evolved derivative of pS4413qbsFH | This study |
| pS4413-qbsFH_A4.F3-I2 | Evolved derivative of pS4413qbsFH | This study |
| pS4413-qbsFGH_A12.F6-I1 | Evolved derivative of pS4413qbsFGH | This study |
| pS4413-qbsFGH_A18.F4-I5 | Evolved derivative of pS4413qbsFGH | This study |
| pGNW2·MetK R281H | Derivative of vector pGNW2 carrying HRs to make the mutation R281H; Km$^R$ | This study |

| | | |
|---|---|---|
| pGNW2·MetK P237T | Derivative of vector pGNW2 carrying HRs to make the mutation P237T; $Km^R$ | This study |

**Supplementary Table 3.** Primers used in this study.

| *Name* | Sequence (5’→3’) | Purpose |
|---|---|---|
| *glyA-I*_g-check_F | TAGTGGATATCCTGGCGTTTGAGCGCCA | Genotyping |
| *glyA-I*_g-check_R | TTTCCTTGCGGAAGCCGATCACTTCGGTT | |
| *glyA-II*_g-check_F | TGATGGAGATTGCCACCTGGCTGCTGCCA | Genotyping |
| *glyA-II*_g-check_R | GCTGCAGATTCTTGCCAGCCTGAAGAAC | |
| *gcv-I*_g-check_F | CCGAACAGGGCAAACAGCAATACCGTAGAC | Genotyping |
| gcv-I_g-check_R | TAGTGCCGATTTCACACCTCCGGCCAGCAG | |
| gcv-II_g-check_F | GATTATTCGAACTGCTCGAGCCCAGC | Genotyping |
| gcv-II_g-check_R | GTTATATCGGCCTGGTGCTGCACAC | |
| pha::M1_g-check_1 | CCTGCGCCTTGTCGAGCTTGC | Genotyping |
| pha::M1_g-check_2 | CCTGCGCCTTGTCGAGCTTGC | |
| pha::M1_g-check_3 | GGCAAGAAGGCCGTCGTGCTCG | |
| pha::M1_g-check_4 | GGCCTCGGTGACGGTGTAGTCG | |
| pha::M1_g-check_5 | CCACGGCTGCAACTCGGTGATCG | |
| pha::M1_g-check_6 | GCAGCCTTGATGGTGTGCTCCAGC | |
| sgRNA_glyA-II_2_F | GCGCGTGCTGGTAAGCCTTGAACTC | Spacer sequence for CRISP/Cas9 counterselection |
| sgRNA_glyAII_2_R | AAACGAGTTCAAGGCTTACCAGCAC | |
| sgRNA_gcv-I_2_F | GCGCGCGAAGCGCATGTAGTAGTGA | Spacer sequence for CRISP/Cas9 counterselection |
| sgRNA_gcvI_2_R | AAACTCACTACTACATGCGCTTCGC | |
| pS4413_BamHI_ F | GGATCCTCTAGAGTCGACCTG | Amplification of vector pS4413 |
| pS4413_KpnI_R | CATATGTTTTTCCTCCTGGTACCGAGCTCGAATTCG | |
| *qbsF*_F | CGAATTCGAGCTCGGTACCAGGAGGAAAAACATATGTGCCCTTGCCCCTACATGC | Amplification of *qbsF*, *qbsH* and *qbsFGH* genes |
| *qbsF*_R | GACGGGGCTCCTGGAAGGTTCAGAGGCTTGAGCGCAGGCTC | |
| *qbsH*_F | CTGAACCTTCCAGGAGCCCCGTC | |

| | | |
|---|---|---|
| qsbH_R | CAGGTCGACTCTAGAGGATCCTTAACCGCTGCGCGGCGC | |
| PS1 | AGGGCGGCGGATTTGTCC | Sequencing pS4413-derived vectors |
| PS2 | GCGGCAACCGAGCGTTC | |
| metK_g-check_1 | ATGAGCGAATACTCCCTTTTCACCTCCGAG | Genotyping |
| metK_g-check_2 | GAGTCGACGATGATCTTGCGGCCAGTCAGG | |
| metK_g-check_3 | CCATGGCTGCGCCCGGATGCCAAGTC | |
| metK_g-check_4 | TTACAGGCCGGCAGCGTCACGCAACG | |
| pGNW_USER_F | AGTCGACCUGCAGGCATGCAAGCTTCT | Amplification of pGNW2 suicide vector for USER cloning |
| pGNW_USER_R | AGGATCUAGAGGATCCCCGGGTACCG | Amplification of pGNW2 suicide vector for USER cloning |
| MetK_USER_R | AGATCCUCGGACGGCTGGCGACATACACC | Amplification of MetK for suicide vector cloning |
| MetK_USER_F | AGGTCGACUGGTGCTCGTCTCCAAGCCCG | Amplification of MetK for suicide vector cloning |
| pGNW_Seq_F | CTTTACACTTTATGCTTCCGG | For colony PCR of pGNW vectors |
| pGNW_Seq_R | TGTAAAACGACGGCCAGT | For colony PCR of pGNW vectors |
| M13R | CAGGAAACAGCTATGAC | Sequencing primer for pGNW vectors |

Synthetic ribosome binding sites (RBS) are underlined, restriction enzyme recognition sites are double underlined.

**Supplementary Table 4.** Whole-genome sequencing methods.

| | DNA extraction | Library preparation | Sequencing |
|---|---|---|---|
| ePUMA | | | |
| A6.F3.I1<br>A8.F3.I1<br>A12.F3.I1<br>A14.F3.I1<br>A16.F3.I1 | Quick-DNA Fungal/Bacteria Miniprep Kit (Zymo Research) | NEBNext Ultra II FS DNA Library Prep Kit for Illumina | Illumina NovaSeq Xplus paired-end 100 bp |
| ePUMA-kyn(WIn) | | | |
| A2.F3.I3<br>A4.F3.I2<br>A6.F3.I1<br>A8.F7.I2<br>A10.F3.I1 | PureLink Genomic DNA Kit (Invitrogen) | Kapa HyperPrep Kit | Illumina NovaSeq Xplus paired-end 100 bp NovaSeq Xplus |
| A2.F3.I1<br>A4.F3.I1<br>A6.F3.I2<br>A8.F7.I1<br>A10.F3.I2 | Quick-DNA Fungal/Bacteria Miniprep Kit (Zymo Research) | SeqWell ExpressPlex™ Library Prep Kit | Illumina NextSeq 2000 paired-end 150 bp |
| ePUMA-Xa(WIn) | | | |
| A12.F6.I1<br>A14.F6.I1<br>A16.F6.I5<br>A18.F4.I5<br>A20.F4.I5 | PureLink Genomic DNA Kit (Invitrogen) | Kapa HyperPrep Kit | Illumina NovaSeq Xplus paired-end 100 bp |
| A12.F6.I2<br>A14.F6.I2<br>A16.F6.I1<br>A18.F4.I1<br>A20.F4.I1 | Quick-DNA Fungal/Bacteria Miniprep Kit (Zymo Research) | SeqWell ExpressPlex™ Library Prep Kit | Illumina NextSeq 2000 paired-end 150 bp |

**Supplementary Table 5.** HR-MS data.

| Compound | Formula | Ion | Calculated | Observed | Δppm |
|---|---|---|---|---|---|
| Xanthommatin | $C_{20}H_{13}N_3O_8$ | $[M+H]^+$ | 424.0776 | 424.0786 | 2.4 |
| Dihydroxanthommatin | $C_{20}H_{15}N_3O_8$ | $[M+H]^+$ | 426.0932 | 426.0941 | 2.1 |
| Decarboxylated xanthommatin | $C_{19}H_{13}N_3O_6$ | $[M+H]^+$ | 380.0878 | 480.0882 | 1.1 |
| Decarboxylated dihydroxanthommatin | $C_{19}H_{15}N_3O_6$ | $[M+H]^+$ | 382.1034 | 382.1039 | 1.3 |

**Supplementary Table 6.** NMR assignments of DC-xanthommatin.

| | **DC-Xa** | |
|---|---|---|
| position | $\delta_C$, type | $\delta_H$ |
| 1 | | |
| 2 | 142.05, CH | 8.44 |
| 3 | 127.33, CH | 7.96 |
| 4 | 119.5, CH | 7.25 |
| 5 | | |
| 6 | | |
| 7 | | |
| 8 | | |
| 9 | 121.38, CH | 7.88 |
| 10 | | |
| 11 | | |
| 12 | | |
| 13 | | |
| 14 | 106.34, CH | 6.82 |
| 15 | 132.97, CH | 7.87 |
| 16 | | |
| 17 | 43.38, $CH_2$ | 3.99, 4.25 |
| 18 | 48.71, CH | 4.46 |
| 19 | | |
